# How can interspecific pollen transfer affect the coevolution and coexistence of two closely related plant species?

**DOI:** 10.1101/2024.09.05.611318

**Authors:** Keiichi Morita, Akira Sasaki, Ryosuke Iritani

## Abstract

Interspecific pollen transfer (IPT), the movement of pollen grains between different plant species by sharing pollinators, incurs costs (fitness reduction) for seed production. IPT thereby reduces the reproductive success of co-flowering plants sharing pollinators, thus preventing their coexistence. However, the impact of IPT on the evolutionary dynamics and evolution-mediated ecological dynamics of sex allocation resource investment to pollen versus ovules) is poorly understood. Here, we investigate the consequences of the female costs incurred by IPT for the co-evolution and coexistence of two plants, by using a mathematical model where two plant species interact with each other via resource competition, pollen movements within and between species, and reduced fertilization due to IPT. The ecological situation we consider here is that an invasive species with female-biased sex allocation immigrates into a habitat of a resident species whose sex allocation is evolutionarily maintained at Fisherian sex allocation (FSA). By using adaptive dynamics theory, we found that regardless of the strength of IPT, natural selection favours the equal allocation to pollen grains and ovules (FSA) for both species. If the mutual impact of IPT on two species is similar in magnitude, we find that the eco-evolutionary dynamics can lead to their stable coexistence. In contrast, when only the invasive species negatively impacts the resident species through IPT, the evolution in invasive species from female-biased sex allocation to FSA causes the extinction of the resident species. Given that local mate competition in small populations is expected to result in female-based sex allocation, our finding suggests that if invasive species are relaxed from local mate competition, they may drive the resident species to extinction. Our study highlights the importance and complexity of the evolution of biased sex allocation driven by IPT to understand the coexistence of closely related plant species.

## 1 Introduction

Transfer of pollen grains commonly occurs between co-flowering plant species with shared pollinators (Morales and Traveset 2008; Moreira-Hernández and Muchhala 2019; Waser 1978). This process is called interspecific pollen transfer (IPT) (Moreira-Hernández and Muchhala 2019). IPT can potentially lower female reproductive success by reducing seed set production. IPT can also reduce male reproductive success by depositing pollen on incompatible stigmas, lowering the chances of successful fertilization (Morales and Traveset 2008; Moreira-Hernández and Muchhala 2019; S. Nishida et al. 2012). Thus, IPT is regarded as a reproductive interference that causes competitive exclusion, hindering coexistence (Kishi and Nakazawa 2013; Levin and Anderson 1970; Moreira-Hernández and Muchhala 2019). Several empirical studies demonstrate that IPT has gained increasing significance among co-flowering species as a result of recent environmental changes. For example, global climate change can shift the range of species distribution, potentially promoting secondary contact, i.e., reunion of allopatrically distributed populations, among co-flowering plants (Parmesan 2006). Increased secondary contact may amplify the impact of IPT on species distribution. Anthropogenic migration may facilitate the establishment of invasive species (Hamann et al. 2021; Suarez and Tsutsui 2008), thereby increasing the exposure of co-flowering species to IPT. Frequent secondary contact and invasion of exotic species may enhance IPT between closely related species that are otherwise geographically isolated (Hamann et al. 2021; Moreira-Hernández and Muchhala 2019; S. Nishida et al. 2012). Therefore, with unprecedented concerns about secondary contact and biological invasions on the rise, it is increasingly important to better understand the impact of IPT on the coexistence of closely related species.

The reduction in reproductive success in both sexes due to IPT not only hinders coexistence but also significantly affects the evolutionary dynamics of reproductive traits (Morales and Traveset 2008; Moreira-Hernández and Muchhala 2019; Waser 1978). IPT promotes the character displacement in the floral structure of closely related co-occurring species (e.g. anther and pollinarium lengths in male function, and style length and stigma size in female function) (Beans 2014; Moreira-Hernández and Muchhala 2019). The relative resource investment in the production of pollen grains over ovules (the P:O ratio, a measure of sex allocation in plants) is an important reproductive trait because the change in P:O ratio influences growth rates of plants by altering the reproductive success of pollen grains and ovules. The sex-specific reduction in reproductive success due to IPT may lead to the evolution of sex allocation to compensate for the loss (Dorken and J. R. Pannell 2009). To our best knowledge, no previous studies addressed how IPT affects the evolution of the P:O ratio, thus leaving it largely unexplored.

The objective of the present work is to explore how IPT influences the joint dynamics of the evolution of sex allocation and coexistence of plant species (see related papers, Rankin, Dieckmann, and Kokko 2011; Zhang and Jiang 1995; Zhang, Lin, and Hanski 2004, for animals). The interplay between the evolution of the P:O ratio and ecological dynamics gains increasing significance because the P:O ratio in isolated populations may become biased, for example, due to local mate competition (Hamilton 1967; West 2009) and other factors (Sato 2004). This increases the likelihood of encounters between closely related species with different P:O ratios upon secondary contact.

Classical theoretical studies on the evolution of sex allocation in plants focused simply on genetic dynamics, assuming a constant population size (D. Charlesworth and B. Charlesworth 1979; Charnov 1988; Holsinger 1991). Several studies have addressed the evolution of reproductive traits in competing species and their impact on coexistence, but they focused on evolutionary rescue driven by rapid evolution of mating cues such as body coloration and demographic oscillation in eco-evolutionary dynamics induced by evolution of prior selfing rate rather than sex allocation or the P:O ratio (Katsuhara et al. 2021; Morita and Yamamichi 2023). Thus, a systematic understanding of the consequences of the evolution of P:O ratios in co-flowering plants for their coexistence via IPT is required to better understand the eco-evolutionary dynamics of plants in nature.

Here, we develop a theoretical model to study the impact of IPT on the coevolution of sex allocation, and examine how the reduction in male and female reproductive success of two closely related species can affect their coexistence. We use adaptive dynamics theory to analyze the eco-evolutionary dynamics (Doebeli 2011; Doebeli and Dieckmann 2000; Hofbauer and Sigmund 1990; Metz, Nisbet, and Geritz 1992), and address the following two questions. First, how does IPT affect the evolution of the P:O ratio? Second, how does an evolutionary change in the P:O ratio affect the coexistence of two closely related plant species? Empirical studies have shown that the strength of IPT is often asymmetrical (Morales and Traveset 2008; Moreira-Hernández and Muchhala 2019); stigmas are sensitive to heterospecific pollen grains, which prevents pollen tube elongation, and this sensitivity varies from one species to another (Etter et al. 2022; Moreira-Hernández and Muchhala 2019). Additionally, animal pollinators have a biased preference for flower morphology (Beans 2014; Bjerknes et al. 2007; Moreira-Hernández and Muchhala 2019; Muchhala 2006; Murcia and Feinsinger 1996). Thus, to evaluate the impact of IPT on the eco-evolutionary dynamics of co-flowering species, we consider the asymmetry in the IPT strength that each species is exposed to. Our paper also aims to demonstrate how asymmetries in IPT strength between species alters eco-evolutionary outcomes.

## 2 Model and Method

### Population dynamics model of competing and pollinator-sharing species

We consider two closely related, hermaphroditic plants that co-flower throughout the year. The two species interact through resource competition and interspecific pollen transfer (IPT). We assume that hybridization does not produce offspring and that selfing does not occur. We also assume that a small population of an invasive species immigrates into a new habitat where a resident species occurs. When the invasive species immigrates, the resident species is assumed to be in equal sex allocation under a large population size, while the invasive species can have a female-biased sex allocation under local mate competition (Hamilton 1967; West 2009) in its native local habitat. The invasive species can also have equal sex allocation if it has a sufficiently large population in its native habitat.

We first have formally constructed models of a fertilization process of female and male gametes in two species that share pollinators and compete for a common resource (D. Charlesworth and B. Charlesworth 1979; Charnov 1988; Holsinger 1991). We consider the mass-action model in which an ovule encounters pollen grains and combine the mass-action models with population dynamics equations of plant individuals. We describe the population dynamics of the two species. The density *N*_*i*_ of species *i* (with *i* = *1, 2*) change with time as

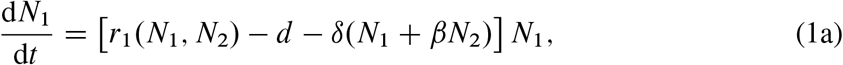

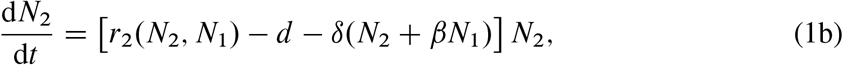

Here, *d* is the natural death rate of both species, *δ* is the coefficient of density-dependent mortality caused by the same species, *δβ* is the corresponding coefficient caused by the other species. Here we have assumed that the interspecific resource competition (IRC) is symmetric between species, that is, intra-specific competition is equally strong in species *1* and *2. r*_*i*_ (*N*_*i*_ ; *N*_*j*_)is the growth rate of species *i*, which may depend on its density *N*_*i*_ as well as the other species’ density *N*_*j*_ (*j* ≠ *i*) through IPT:

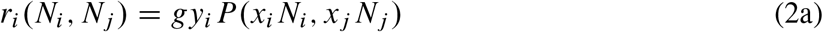

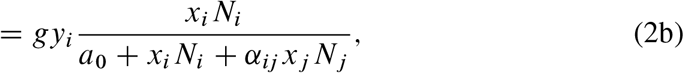

where *x*_*i*_ and *y*_*i*_, respectively, is the per capita pollen and ovule production by an individual of species *i, P*(*x*_*i*_ *N*_*i*_, *x*_*j*_ *N*_*j*_) = *x*_*i*_ *N*_*i*_ *=*(*a*_0_ + *x*_*i*_ *N*_*i*_ + *α*_*ij*_ *x*_*j*_ *N*_*j*_) is the probability that each ovule encounters conspecific pollen, which is a function of the total pollen production, *x*_*i*_ *N*_*i*_, of the same species and that of the other species, *x*_*j*_ *N*_*j*_. The strength of IPT is measured by the relative rate *α*_*ij*_ at which an ovule is fertilized by heterospecific pollen, rather than conspecific one. The parameter *a*_0_ in the denominator of (Eq. 2b) measures how the fertilization rate is limited by the total pollen production. If *a*_0_ = *0*, then there is no pollen limitation (all ovules can encounter pollen). With a nonzero *a*_0_, the fewer the total pollen production, the greater the probability that an ovule fails to encounter pollen. If *x*_*i*_ *N*_*i*_ is equal to *a*_0_ in the absence of the competing species *j*, the growth rate is *gy*_*i*_ *=2*, which is a half value of the maximum growth rate. In this sense, *a*_0_ is half saturation pollen density in the absence of the competing species. For the derivation of this form of *P* from biological processes, see Supporting information. The other factor involved in the growth rate *r*_*i*_ is the germination rate *g* (the probability that a fertilized ovule will form seed and germinate). An overview of the model is given in Fig. 1.

**Figure 1.**
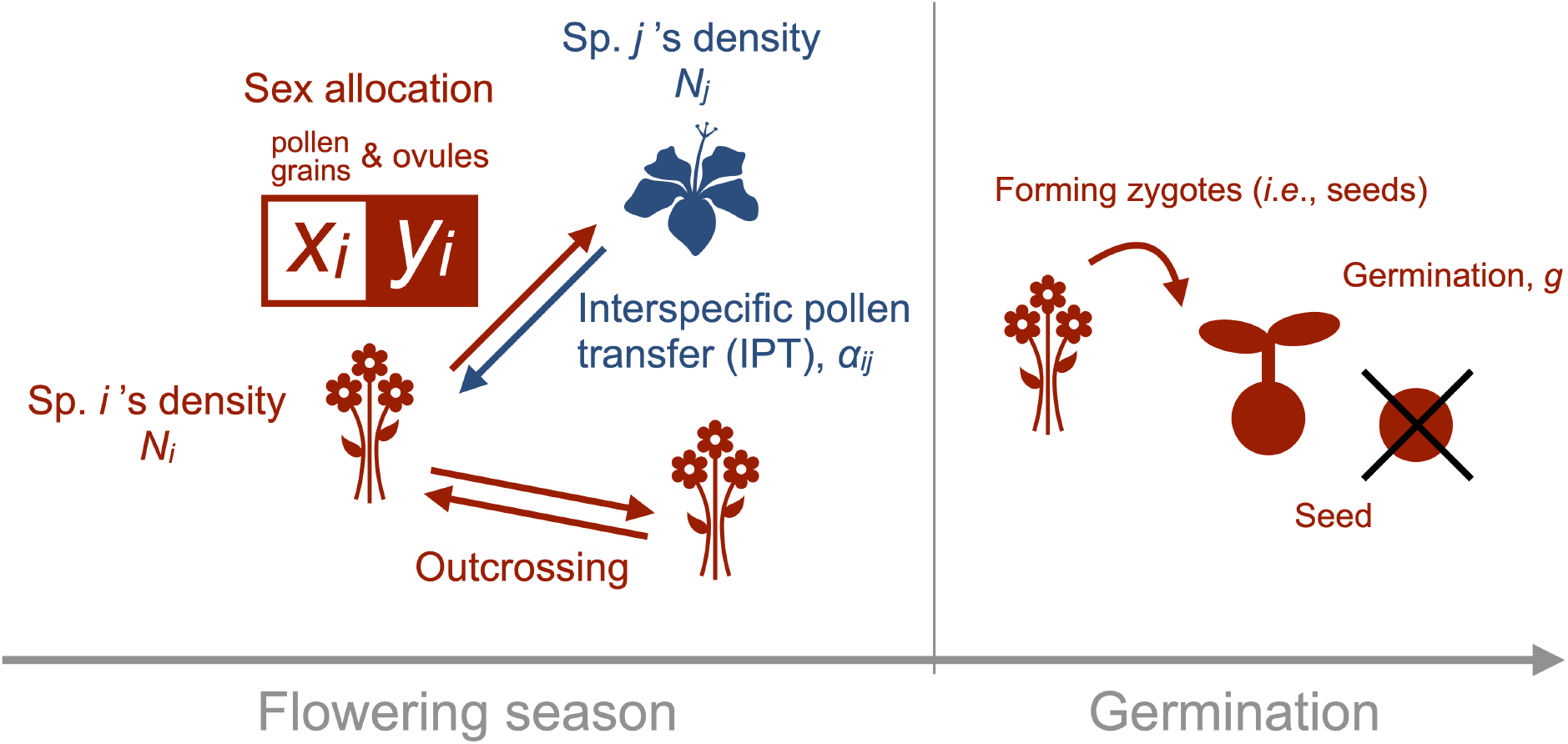
Model overview: We incorporate sex allocation into a population dynamics model of co-flowering plants, species 1 and 2 with a density of adult flowers, *N*_1_ and *N*_2_, respectively. The flowers of a plant of species *i* (*i* = 1, 2) produce the number *x*_*i*_ of pollen grains and *y*_*i*_ of ovules *x*_*i*_ and *y*_*i*_, respectively. The ratio of *x*_*i*_ to *y*_*i*_ represents resource allocation to male and female functions (sex allocation). Species *i* receives conspecific pollen grains at a constant rate and heterospecific pollen grains of species *j* at a constant rate weighted by a parameter, *α*_*ij*_ (*i*≠*j*). As *α*_*ij*_ becomes large, more ovules encounter heterospecific pollen grains, reducing the population growth rate of species *i*. Therefore, *α*_*ij*_ is one of important parameters indicating the strength of IPT. When ovules encounter pollen grains, zygotes are formed and the zygotes germinate at a rate, *g*.

Increasing per capita ovule production, *y*_*i*_, increases the growth rate (Eq. 2a). Increasing per capita pollen production, *x*_*i*_, also contributes to increasing the growth rate by increasing the probability *P* of an ovule encountering conspecific pollen grains (Eq. 2). In addition, increasing the pollen production of one species will decrease the growth rate of the competing species by preventing their ovules from encountering conspecific pollen grains (this effect is seen in Eq. 2b by increasing *x*_*j*_). One of the objectives of the present paper is to examine how the reduction in female reproductive success induced by mating with heterospecific pollen grains affects the evolution of sex allocation. This might suggest that producing more pollen grains is beneficial under the intraspecific competition with co-flowering (pollinator sharing) species. However, we can easily show that both species evolve towards equal sex allocation even when IPT occurs. This is because, while male-biased sex allocation of one individual benefits the other individuals of the same species by reducing the fecundity of competing species through IPT, other individuals who allocate equally to pollen and ovules can have more benefit than the individuals with male-biased sex allocation. This will be discussed in detail in Results.

As the per capita total resource allocation to male and female functions is limited (Campbell 2000; D. Charlesworth and B. Charlesworth 1981; Charnov 2020), we assume a linear trade-off between investment to produce pollen grains, *x*_*i*_, and ovules, *y*_*i*_, in total resource allocation, *R*:

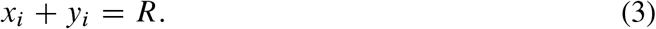

To simplify the analysis of ecological dynamics, we assume that there is approximately no pollen limitation (*a*_0_ = *0*) (Supporting information for details). Under these assumptions, IPT leads to a frequency-dependent process of reproductive interference (as in, for example, Kishi and Nakazawa 2013; Kuno 1992):

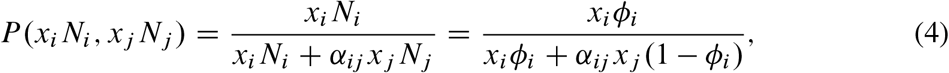

where *ϕ*_*i*_ = *N*_*i*_ *=*(*N*_*i*_ + *N*_*j*_) is the frequency of species *i* among co-flowering species. The numerical analysis shows that the dynamics with *a*_0_ = 0 gives qualitatively similar result to that with small but positive *a*_0_ (Fig. S5a-c).

We first summarize our simplifying assumptions before addressing the results. To focus on the effect of IPT on evolutionary and ecological consequences, we assume that the two species have the same parameters except for the strength of IPT,*α*_*ij*_, and the allocation to pollen grains and ovules, *x*_*i*_ and *y*_*i*_. That is, two species may differ in their investments, *x*_*i*_ and *y*_*i*_, to pollen and ovules, and may have different impacts of IPT to the other, but are otherwise equal: they have the same values of the germination fraction, mortality rate, total investment in the production of pollen grains and ovules, and have the same coefficients of intra- and interspecific resource competition. These are the additional assumptions other than pollen limitation is negligible (*a*_0_ = 0). All variables and parameters are defined in Table 1.

**Table 1:** Symbols and Their Definitions.

| Symbols | Definition | Default values |
| --- | --- | --- |
| $N_i$ | Density of species $i$ | Dynamic variable |
| $R$ | The total resource a plant invests in the production of pollen grains and ovules | 20 |
| $x_i$ | Pollen production of a plant | Evolutionary trait |
| $y_i$ | Ovule production of a plant | $R - x_i$ |
| $g$ | Germination fraction | 0.83 |
| $d$ | Natural mortality rate | 0.1 |
| $\delta$ | Coefficient of density-dependent mortality | 0.01 |
| $a_0$ | Half saturation pollen density in the absence of competing species | 0 |
| $\alpha_{ij}$ | The rate at which a heterospecific pollen grain is transferred to an ovule (relative to that of a conspecific pollen grain) | $0 \leq \alpha_{ij} \leq 1$ |
| $\beta$ | The strength of interspecific resource competition relative to that of intraspecific competition | $0 \leq \beta \leq 1$ |

Under these assumptions, we can carry out a conventional variable-transform (Otto and Day 2007) so that otherwise two-species dynamics can be converted to the dynamics of a ratio of densities of two species: *ρ* = *N*_*1*_*/N*_*2*_. By analyzing the dynamics of *ρ*, we examine how the coexistence condition changes by changing the sex allocation parameter *x*_*i*_ of invasive species (Figs. S1a, 2a, 3a). The derivation of the dynamics of *ρ* is detailed in Supporting information.

### 2.1 Adaptive dynamics theory

#### Invasion fitness

To investigate long-term co-evolutionary dynamics of sex allocation, we use adaptive dynamics (Doebeli 2011; Doebeli and Dieckmann 2000; Hofbauer and Sigmund 1990; Metz, Nisbet, and Geritz 1992). Specifically, we first consider a phenotypically monomorphic population of a wild type and then introduce a rare mutant that has a slightly different phenotype, 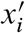, attempting to invade the population of the wild type. That is, we assume that natural selection is weak because phenotypic difference between the mutant and wild type is small: 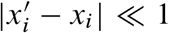. As such, evolution occurs much more slowly than population dynamics, and the monomorphic population (Eq. 1) reaches a demographic equilibrium, 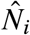 and 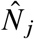 with the phenotype of the wild type, *x*_*i*_ for each species. A rare mutant in species *i* can invade the population of the wild type if its invasion fitness is positive and fails to invade if it is negative. The invasion fitness of a mutant with phenotype 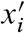 in the population of the wild type of species *i* having phenotype *x*_*i*_ and that of species *j* having phenotype *x*_*j*_ is

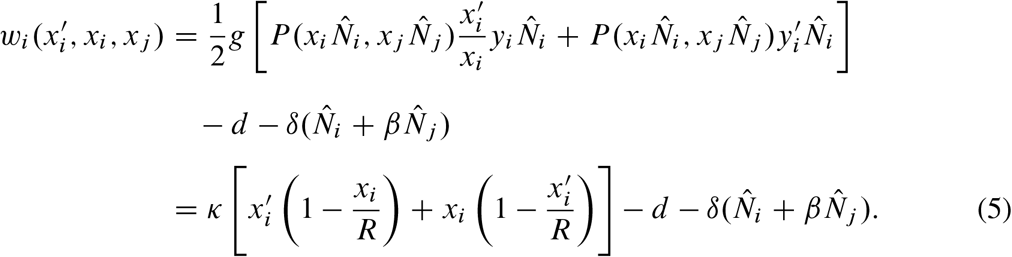

See Supporting information for derivation. The parameter 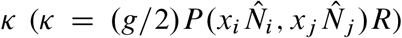 is a positive factor that is independent of the mutant trait 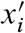. The first term of the mutant’s invasion fitness in Eq. (5) is given by the sum of the reproductive successes through pollen grains and ovules, while the second and third terms respectively represent the natural mortality and density-dependent mortality due to resource competition.

#### Convergence and evolutionary stability

To investigate the direction of sex allocation evolution, we calculate the selection gradient, *D*_*i*_ (*x*_*i*_, *x*_*j*)_, for the allocation *x*_*i*_ of species *i* to male function, given that the species *j* allocate *x*_*j*_ to male function, from the invasion fitness *w*_*i*_ of the mutant of species *i* :

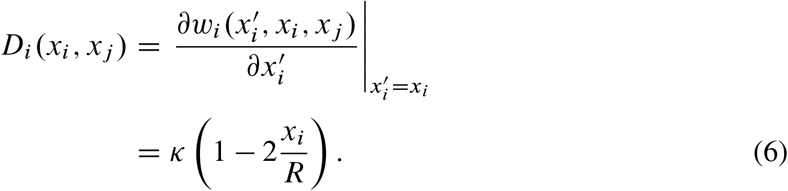

Solving *D*_1_(*x*_1_, *x*_2_) = *D*_2_(*x*_2_, *x*_1_) = 0 for *x*_1_ and *x*_2_, we have an evolutionary equilibrium 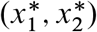 at which natural selection ceases, called evolutionarily singular strategy. Furthermore, the sign of selection gradient determines the direction of natural selection acting upon the allocation strategy. If *x*_*i*_ */ R* = *0:5*, the selection gradient is zero. In the case where trait *x*_*j*_ of species *j* is fixed, natural selection in species *i* favors a mutant that has a higher allocation to males (or females, respectively) if the selection gradient *D*_*i*_ (*x*_*i*_ ; *x*_*j*_) is positive (or negative, respectively). In the other case where both species *i* and *j* evolve, we find whether or not evolution converges into the singular point by examining real parts of eigenvalues in the following Jacobian matrix (Dieckmann 1995; Dieckmann and Law 1996; Doebeli and Dieckmann 2000):

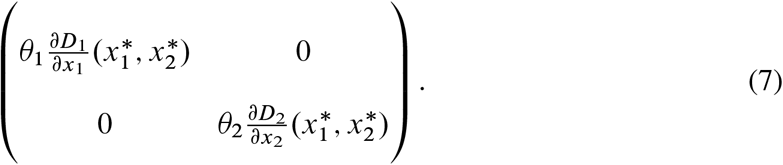

where *θ*_*i*_ is the constant evolution rate determined by the mutation rate and population density of each species at equilibrium. The Jacobian matrix is diagonal in this system, which indicates that the sex allocation strategies of two species evolve independently. When all the real parts of the eigenvalues of Eq. (7) are negative, the biological system approaches the evolutionarily singular point (i.e. the singular point is “convergence stable”).

The selection gradient does not tell whether or not the singular strategy is “evolutionarily stable,” which is determined by the fitness curvature (Dieckmann 1995; Dieckmann and Law 1996; Doebeli and Dieckmann 2000; Smith 1982): a singular strategy is evolutionarily stable (Smith 1982) (i.e. it refuges the invasion of any other nearby mutants) if the curvature of invasion fitness evaluated at the singular strategy is negative:

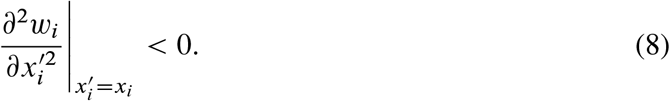

An evolutionarily singular strategy is neutrally evolutionarily stable if the fitness curvature is zero.

## 3 Results

We raised two questions in the Introduction: (1) How does interspecific pollen transfer (IPT) affect the evolution of P:O ratio? (2) How does an evolutionary shift in P:O ratios affect the coexistence of two closely related plant species? We answer the first question by showing that the P:O ratio evolves to equal allocation to pollen grains and ovules regardless of the IPT between species. To answer the second question, we first show how the coexistence of two species competing through IPT is affected by the sex allocation difference between them. We next examine how an evolutionary shift in sex allocation alters the outcomes of ecological dynamics. We here focus on the case where two species compete only through IPT (i.e. the interspecific resource competition (IRC) is negligible). We later show that the result remains qualitatively the same when IRC exists but is weak.

We found that the results of our model are best summarized by classifying biological situations into three categories differing in relative strengths *α*_*ij*_ of IPT between competing species: (i) symmetric IPT in which the IPT from species *i* to *j* is equally strong as that from *j* to *i* (*α*_12_ = *α*_21_) (ii) IPT from an invasive species 2 is much stronger than IPT from a resident species 1 (*α*_12_ » *α*_21_) (iii) IPT from the resident species 1 is much stronger than IPT from the invasive species2 (*α*_21_ » *α*_12_).

### 3.1 How does IPT affect the evolution of P:O ratio?

As explained in Model and Method, the joint evolutionary equilibrium for sex allocation strategies of species 1 and 2 are obtained from the condition that selection gradients for sex allocation strategies vanish in both species, *D*_1_(*x*_1_, *x*_2_) = *D*_2_(*x*_*2*_, *x*_1_) = 0, which yields 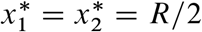. Under this equilibrium, both species invest equally in male and female functions, 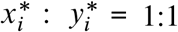 (Fisherian sex allocation, FSA) (Charnov 2020; Fisher and Bennett 1999; West 2009). This equilibrium 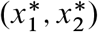 is always convergence stable (Leimar 2009) as both eigenvalues of the Jacobian matrix (7) are negative (Eq. (S24) of Supporting information). We can also show that the equilibrium is neutrally evolutionarily stable (i.e. 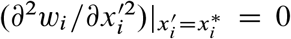) for both species i=1,2) as in the classical Fisherian sex ratio theory. Therefore, the model analysis predicts that the sex allocation of two species coevolves towards the FSA. This result is consistent with numerous theoretical studies that show that selection leads to an equal investment in male and female functions (Fisher and Bennett 1999) regardless of differential mortality rates after parental investment (e.g. Pirrie and Ashby 2021) and sex-biased mortality in adulthood (Kokko and Jennions 2008; West 2009).

### 3.2 How does an evolutionary shift in P:O ratios affect the coexistence of two closely related plant species?

Assuming that there is no interspecific resource competition (IRC) (*β* = *0*), we obtained three different outcomes corresponding to three cases differing in the relative strengths of IPT between species. Similar results are obtained according to the numerical analyses for the case of weak IRC (*β* is positive but small). Under these conditions, the outcomes of eco-evolutionary dynamics are drastically different depending on the following two factors: the strength of IPT from the invasive species to the resident species relative to IPT from the resident species to invasive species, and the initial sex allocation of the invasive species. As for the relative strength of IPT, we consider three cases: (i) the strength of IPT from the invasive species 2 to the resident species 1 is nearly the same as that of the opposite direction (*α*_12_ *α*_21_) or if neither strength of IPT is not too large (the shaded area in Fig. 2a). (ii) the strength of IPT from invasive species 2 is much stronger than that from the resident species 1 (*α*_12_≈ *α*_21_) the right white area in Fig. 2a), and (iii) the strength of IPT from resident species 1 is much stronger than that from the invasive species 2 (*α*_12_ ≪ *α*_21_) the upper white area in Fig. 2a). As we mentioned in Model and Method, the resident species is assumed to have equal sex allocation (FSA) before the immigration of invasive species. As we have shown in the previous section, the sex allocation of each species evolves independently to FSA even under the reproductive interference via IPT from the other species. Therefore, the resident species stays in FSA throughout the eco-evolutionary dynamics. On the other hand, the invasive species can initially have a biased sex allocation, under, for example, local mate competition at their native habitat (Hamilton 1967). Now we summarized the results for the cases (i)-(iii).

**Figure 2.**
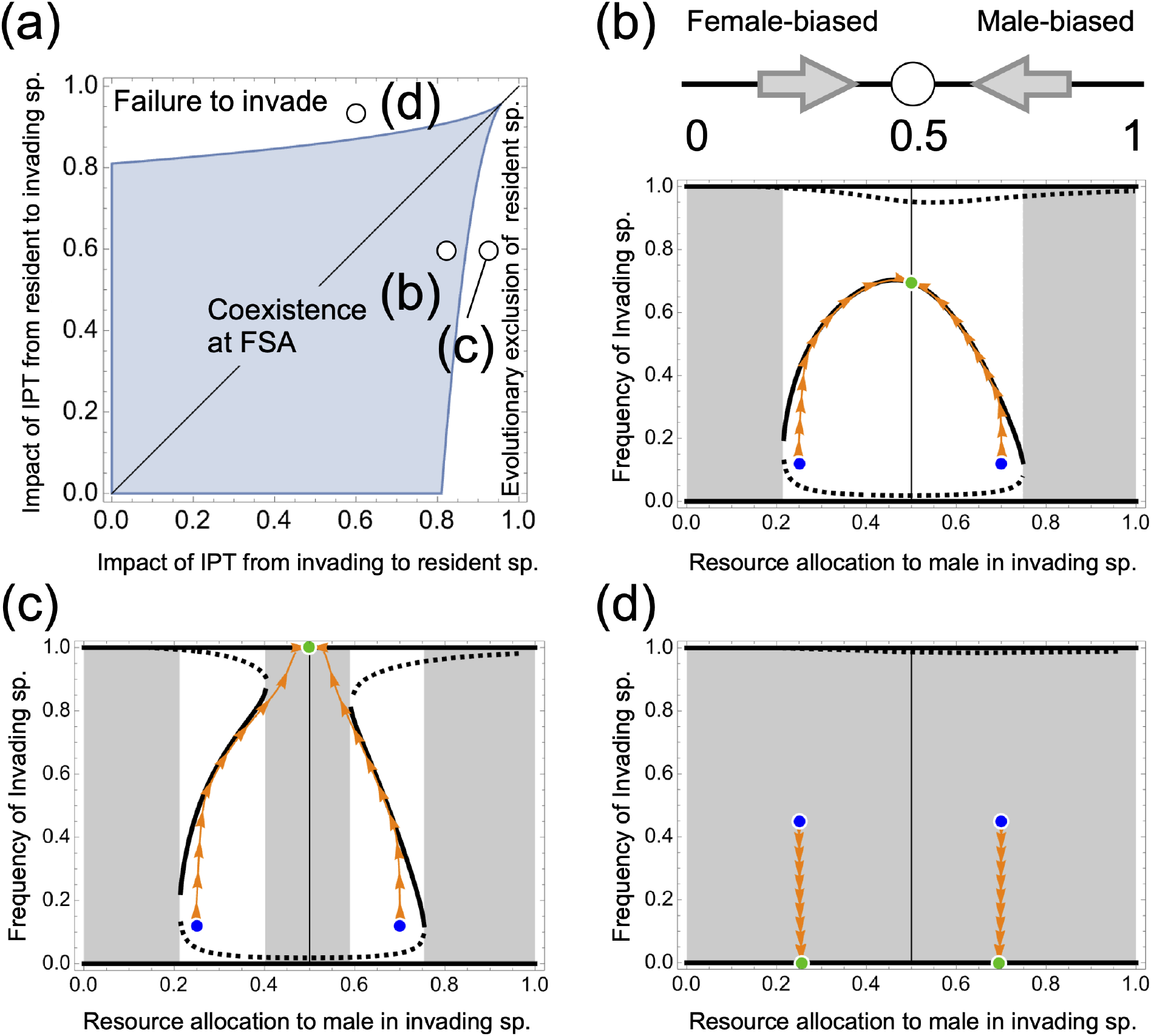
The relationship between the strength of interspecific pollen transfer (IPT) and evolutionary exclusion: (a) The phase diagram classifying the eco-evolutionary outcomes into three types. The horizontal axis represents the strength of IPT from an invasive to a resident species, while the vertical axis represents the strength of IPT in the opposite direction. (b, c, d) The trajectories of eco-evolutionary dynamics (lines with arrows) plotted against the sex allocation of invasive species (allocation to males, *x*_*i*_ ; horizontal axis) and the relative density, *ρ*= *N*_1_*/N*_1_ + *N*_2_), of invasive species in the two-species community (vertical axis). In a white region, there is a stable internal ecological equilibrium (solid thick line) together with two unstable equilibria (thick dashed lines). In the shaded region, there is no internal ecological equilibrium, indicating that either the invasive or resident species always deem to go extinct. The sex allocation of the resident species (not shown) remains at FSA if started from FSA (see text). White open cirlces show the values of *α*_12_ and α_21_ used in panels (b), (c), and (d). (b: Top) (b) (Top diagrams) Even under IPT, initially biased sex allocation of invasive species always evolves to FSA, regardless of parameters. (b: Bottom) When IPT is nearly symmetric between species, the immigration of invasive species with possibly a biased sex allocation will end up with the stable coexistence of two species with equal sex allocation in each, provided that initial sex allocation is only intermediately biased (white region). In contrast, if the invasive species has an extremely biased sex allocation, the invasive species goes extinct (gray region). (c) If the strength of IPT from the invading species is stronger than that from the resident species, the evolution of the invading species results in the extinction of the resident species. (d) If the strength of IPT from the resident species is stronger than that from the invading species, the invading species always goes extinct. The strengths of IPT are (b) *α*_12_ = 0.8, *α*_21_ = 0.6, (c) *α*_12_ = 0.9, *α*_21_ = 0.6, (d) *α*_12_ = 0.6, *α*_21_ = 0.9. The other parameters are in Table 1.

(i) If neither IPT from the invasive species nor that from the resident species is strong, or if they are nearly symmetrical (as shown in the shaded area in Fig. 2a), two co-flowering species can stably coexist despite reproductive interference by IPT, even if the invasive species initially has a biased sex allocation. The trajectory of eco-evolutionary dynamics (orange arrows in Fig. 2b) shows that the allocation of the invasive species to males (the horizontal axis) evolves to FSA (*x*_2_ = 0.5). Meanwhile, the frequency *ϕ*_2_ of the invasive species (the vertical axis) approaches an internal equilibrium. This implies that the invasive species stably coexists with the resident species. At this eco-evolutionary equilibrium, both species invest equally in males and females, and their pollen grains continue to interfere with each other’s reproduction.

Two additional points should be noted. First, if the invasive species initially has an extremely biased sex allocation (gray area in Fig. 2b), its cost in growth rate is so great and the invasive species is driven to extinction because there are too few pollen grains to encounter ovules and vice versa. Second, if the initial population of invasive species is too small (the frequency below the dashed line near the bottom in Fig. 2b), the invasive species fails to invade irrespective of its initial sex allocation. This is due to the bistability of two marginal equilibria where only one species can exist, caused by the majority advantage in reproductive interference (Kishi and Nakazawa 2013; Kuno 1992).

(ii-a) If the strength of IPT from the invasive species is much stronger than that from the resident species (the right white area in Fig. 2a; *α*_12_ » *α*_21_), the invasive species drives the resident species to go extinct if the invasive species initially has nearly equal sex allocation (a gray area in Fig. 2c).

(ii-b) More complex eco-evolutionary interaction arises in the case *α*_12_ » *α*_21_ if invasive species initially has an intermediately biased sex allocation. Then, the invasive species and the resident species temporarily coexist at quasi-equilibrium (thick curve in Fig. 2c). However, the sex allocation of the invasive species gradually evolves towards FSA, pressing the temporarily coexisting population until it falls into the gray area for the extinction of the resident. After the population falls into the gray area, it shows the same behavior as described in (ii-a): the invasive species drives the resident species to extinction. We refer to such extinctions by the evolution of a competitor species as “evolution-mediated competitive exclusion.”

The same additional notes apply as in the case (i). If the initial sex allocation of the invasive species is extremely biased, the invasive species fails to invade due to too large cost for sex allocation bias. If the initial population of invasive species is too small in the white area of Fig. 2c, the invasive species go extinct due to the majority advantage for the resident species in reproductive interference (Kishi and Nakazawa 2013; Kuno 1992).

Finally, (iii) if the strength of IPT from the resident species is much stronger than that from the invasive species (*α*_21_ » *α*_12_; the upper white area in Fig. 2a), the invasive species with any sex allocation cannot establish itself because the invasive species is suffered too strongly from the cost incurred by IPT from the resident species (Fig. 2d).

We now focus on the mechanisms of ecological coexistence (the white areas of Fig. 2b and 2c). If the invasive species has male-biased allocation, it hinders more the reproduction of the resident species than the resident species hinders the reproduction of the invasive species. This is because the invasive species sends more pollen grains to the ovules of resident species than vice versa. However, the over-production of pollen grains by the invasive species comes at the cost of reduced ovule production. The balance between the more negative impact of IPT from invasive species on the resident species and the lower ovule productivity of the invasive species allows them to coexist. If the invasive species has a female-biased allocation, it suffers more from IPT caused by the resident species. However, this disadvantage is offset by its higher ovule production compared to the resident, which facilitates coexistence.

### 3.3 Robustness of the results

#### 3.3.1 What Happens When Initial Sex Allocation of the Resident Species Deviates from FSA

So far, we have assumed that the resident species has equal sex allocation (FSA) when the invasive species immigrates. Essentially the same results follow as we have shown in Fig. 2 even when the initial sex allocation of resident species deviates from FSA (Figure S1a for case (i) and S1d for case (ii) in the previous section).

This is because sex allocation in both invasive and resident species evolves to FSA independently with each other and independently of ecological conditions including reproductive interference through IPT. Conversely, the ecological dynamics of these species are largely impacted by to what extent each species invests in males and females. As a result, the eco-evolutionary outcome is determined by how the evolutionary trajectory moves through areas of qualitatively different ecological consequences (the white, gray, and black regions in Fig. S1a and S1d). The robustness follows from the fact that the regions and the order in which evolutionary trajectories pass through when the resident species’ sex allocation is restricted to FSA is the same as the regions and the order in which they pass through when such restriction is relaxed (for example, Fig. S1b, c for case (i) and Fig. S1e, f for case (ii)).

#### 3.3.2 What Happens When Two Species Also Compete for Resources

For the sake of mathematical simplicity, we have assumed in the previous section that two species compete only through the IPT (i.e. there is no interspecific resource competition: *β* = 0). The analysis for a positive but small *β* is shown in Figure S2. Fig. S2e-h (with *β* = 0.1) correspond to Fig. 2a-d (with *β* = 0), which are qualitatively similar. There are, however, two notable differences between the case *β* = 0 and *β* = 0.1. First, the region for the presence of ecoevolutionary equilibrium in which both species coexist at FSA (the shaded region in Fig. S2e) is narrower in the case *β* = 0:1 than in the case *β*= 0. Additional competition for resources makes the stable coexistence at evolutionary equilibrium difficult. Second, the asymmetry between the effect of male- and female-biased sex allocation of invasive species is amplified with *β* = 0:1 than with *β* = 0 (see, for example, the thick line, the stable internal equilibrium of ecological dynamics, in Fig. S2f-g.

When the parameters *β*, the strength of interspecific competition (IRC), and *α*, the strength of interspecific pollen transfer (IPT) are varied more widely, the model shows more complex behavior (Fig. S3). We see this by focusing on the two most interesting results of our model: ‘evolution-mediated competitive exclusion’, where the evolution of sex allocation in invasive species drives the resident species to extinction, and the coexistence of two species with FSA. As full search for parameters is too complicated, we here restricted ourselves to the case of symmetric IPT: *α*_12_ = *α*_21_ = *α*. The results are summarized in Fig. S3a. First, the region for which the evolution-mediated competitive exclusion can occur is found in a wide range of *β* and *α*, though the region (shown in black in Fig. S3a) is narrow in *β* -*α* space and eco-evolutionary dynamics is described in Fig. S3e. For the evolution-mediated competitive exclusion to occur, the strength of IPT and IRC must be negatively correlated: if IPC is strong, IRC must be weak, and vice versa. Second, the region for which the coexistence of two species with FSA occurs is found in a broader area in *β* -*α*space (shown in gray in Fig. S3a), surrounding the region for which the evolution-mediated competitive exclusion occurs and eco-evolutionary dynamics is described in Fig. S3b-d, f. Third, if both IPT and IRC are strong, competitive exclusion occurs without invoking evolutionary changes (shown in white in Fig. S3a) and eco-evolutionary dynamics is described in Fig. S3g.

Our results are robust for the changes in other parameter values including half saturation pollen density in the absence of competing species, *a*_0_, natural mortality of adults, *d*, and the total resource invested to produce pollen grains and ovules, *R* (Fig. S5) although large changes in parameter values may result in qualitatively different outcomes (Supporting information).

## 4 Discussion

Through rigorous theoretical analysis, our research has uncovered two significant findings with important implications for the co-evolutionary dynamics of sex allocation in a two-species plant community and community dynamics. First, the sex allocation of each of co-flowering species under interspecific pollen transfer (IPT) evolves towards the Fisherian sex allocation (FSA). The second finding illuminates how two interdependent processes of evolution of sex allocation and population dynamics ultimately shape the outcome of species coexistence. We have found that the evolution of sex allocation in invasive species may drive resident species to extinction, a process we refer to as ‘evolution-mediated competitive exclusion.’ Using mass-action models, we have formally modeled a fertilization process of female and male gametes (D. Charlesworth and B. Charlesworth 1979; Charnov 1988; Holsinger 1991), and as such, we have described the effect of IPT on the reproductive success of the species. Building upon the formulation, we have further constructed and analyzed co-evolutionary dynamical models of sex allocation, thereby revealing the consequences of the evolution of sex allocation for the coexistence of two closely related plant species.

Our results show that evolution-mediated competitive exclusion may occur if interspecific resource competition (IRC) is weak and IPT from invasive species is stronger than from resident species (Fig. 2a, Fig. S2). Although both IRC and IPT commonly occur in nature, we find that IPT results in evolution-mediated competitive exclusion regardless of the strength of IRC. As in our assumption of evolution-mediated competitive exclusion, the asymmetric effect of IPT may be common in reality (Morales and Traveset 2008; Moreira-Hernández and Muchhala 2019), for instance, due to variations in pollinator preference and floral morphology (Bjerknes et al. 2007; Etter et al. 2022; Moreira-Hernández and Muchhala 2019; Muchhala 2006; Murcia and Feinsinger 1996; Natalis and Wesselingh 2012). When an invasive species has a less conspicuous flowering color or stamens positioned far from the pistils, the ovules of the resident species receive more pollen from other species than those of the invasive species do. Our model suggests that the evolution of sex allocation in invasive species, when associated with asymmetric IPT, may lead to the extinction of resident species. Additionally, our results indicate that even if IRC is strong and IPT is symmetric and weak, evolution-mediated competitive exclusion can occur, due to the combined negative impacts of IPT and IRC (Fig. S3). We also found that evolution-mediated competitive exclusion is likely to occur when the strengths of IPT and IRC are negatively associated (e.g., when IPT is strong and IRC is weak, or vice versa). These predictions, which may be relevant to a wide range of biological systems in nature, are testable in experimental settings. Our findings have practical implications for the management and conservation of plant species facing environmental changes and species migration followed by human-mediated species translocation.

The evolution-mediated competitive exclusion is driven by the evolution of sex allocation in invasive species from female-biased to equal allocation (FSA). Female-biased sex allocation may occur due to local mate competition in spatially structured populations (Hamilton 1967; West 2009). When individuals of the invasive species immigrate from their native populations in a fragmented habitat to a large habitat where the resident species resides, natural selection tends to favor less female-biased sex allocation or even FSA in immigrated invasive species, because individuals more likely encounter non-relative mating partners than in their native habitat.

Empirical studies have observed that invasion of closely related species reduces the population of native species due to IPT (Chittka and Schürkens 2001; Kephart 1983; Matsumoto, Takakura, and T. Nishida 2011). Other studies reported geographic variation in sex allocationand other floral traits such as flower size and seed traits between hermaphroditic populations in the introduced and native habitats (Boheemen, Atwater, and Hodgins 2019; Fenollosa, Jené, and Munné-Bosch 2021; Goldman and Willson 1986; Montague, Barrett, and Eckert 2008; J. Pannell et al. 2014). These lines of research indicate that the sex allocation in plants may be subject to natural selection associated with invasion processes. As such, our results can be validated by field studies comparing the sex allocation of invasive species in their native and introduced habitats.

Global warming and human-mediated species translocation have recently caused a shift in species distribution and a long-distant migration (Hamann et al. 2021; Parmesan 2006; Suarez and Tsutsui 2008). This increases the likelihood for a resident species to encounter a closely related species. This suggests that a drastic shuffling in community structure including evolution-mediated competitive exclusion proposed in this paper would occur frequently in the Anthropocene.

Our study shows that evolutionary changes in interacting species after the secondary contact can have a significant impact on their ecological dynamics, through reproductive interference resulting from interspecific pollen transfer (IPT) between co-flowering plant species. Recent research highlights the importance of IPT in elucidating the relationship between adaptive evolution and the coexistence of species (Gómez-Llano et al. 2021; Svensson 2019). Morita and Yamamichi (2023) found that the rapid evolution of reproductive traits such as male ornaments in rare species facilitates population-recovery as a result of reproductive character displacement (Morita and Yamamichi 2023). Katsuhara et al. (2021) showed that IPT-driven evolution of prior, autonomous selfing results in long-term coexistence with oscillations (Katsuhara et al. 2021). These studies focused on coexistence due to ‘evolutionary rescue’ in which the reduction in the strength of reproductive interference and IPT due to character displacement cause the recovery of populations of rare species, and discussed the conditions for evolutionary rescue by focusing on the effect of the magnitude of additive genetic variance (Morita and Yamamichi 2023), and the nature of the breeding system (Katsuhara et al. 2021). In contrast, our paper focuses on the evolution of sex allocation under IPT and shows that the evolution from sex biased allocation to FSA in one species results in competitive exclusion of the other non-evolving species (evolution-mediated competitive exclusion). Evolution-mediated competitive exclusion occurs because the evolutionary convergence from biased sex allocation to FSA in evolving invasive species strengthens the negative effect of IPT on the growth rate of non-evolving resident species at FSA, while the models of Morita and Yamamichi (2023) and Katsuhara et al. (2021) consider the evolutionary divergence of reproductive traits that weaken, rather than strengthen, reproductive interference and IPT. Therefore, the evolutionary convergence of the reproductive traits drives competitive exclusion, while the evolutionary divergence promotes coexistence.

Our paper considers a situation where pollen grains are abundant, indicating no male cost incurred by IPT. Many plants produce sufficiently abundant pollen grains, making the male cost incurred by IPT negligible (Cruden 2000; Moreira-Hernández and Muchhala 2019). In some case, we cannot ignore the male cost. For instance, fewer pollen grains are produced in self-compatible plants than in self-incompatible plants (Cruden 2000). Also, many pollen grains are lost before they encounter with conspecific ovules due to interspecific pollen competition and the low efficiency of pollination (Cunha et al. 2022; Harder and Johnson 2023; Moreira-Hernández and Muchhala 2019). However, previous theoretical works showed that equal sex allocation evolves in the presence of the male cost in a large population with selfing (Bochynek and Burd 2024; Charnov 2020; Crowley, Harris, and Korn 2017; Queller 1984). Therefore, even if we assume the male cost, our results will be qualitatively same, provided that we consider the ecoevolutionary dynamics in a large population. Our model can be applied to various situations in the presence of the male cost. Note that in small population, the theoretical works suggested that the male cost results in evolution of biased sex allocation (Bochynek and Burd 2024; Crowley, Harris, and Korn 2017). Empirical studies reported that if populations are located in the high latitude where pollinators are scarce and pollination is inefficient, the populations have more biased sex allocation than if pollination is efficient (Cunha et al. 2022; Harder and Johnson 2023; Moreira-Hernández and Muchhala 2019). Thus, eco-evolutionary outcomes will be changed if we consider two plant species with a small population.

Although we did not consider selfing, biological invasion of plants is often associated with natural selection on selfing, a crucial factor to establishment success (Cutter 2008, 2019). “Selfing syndrome” is one of the examples indicating the importance of the evolution of selfing in invasive species (Cutter 2008, 2019). In selfing syndrome, there are evolutionary transitions from outcrossing to selfing after immigration in numerous plants (Cutter 2008, 2019). A previous theoretical study suggested that the evolution of prior, autonomous selfing can reduce the IPT-induced reduction in growth rates and thus promote coexistence (Katsuhara et al. 2021). Also, previous studies suggested that sex allocation (pollen:ovule ratio; P:O ratio) is more female-biased in selfing hermaphrodites than self-incompatible, outcrossing plants (Charnov 2020). Therefore, our prediction generates the hypothesis that the evolution of selfing in invasive species may maintain female-biased sex allocation and prevent evolution-mediated competitive exclusion. Incorporating selfing into our model will help formulate this hypothesis, and may elucidate how diverse mating systems in plants can allow for coexistence.

Flowering time are important characteristics of mating systems in plants in addition to the reproductive structures (Baker, Burd, and Climie 2005; Eckhart 1993; Klinkhamer and De Jong 1987). The reproductive structures is sometimes linked to flowering time genetically. For example, Eckhart showed that high seed production is associated with early flowering in experimental populations of *Phacelia lineans* (Eckhart 1993), indicating that the evolution of sex allocation, which changes the ovule production level, can strongly influence flowering time. This further suggests that evolutionary shift in sex allocation may lead to speciation via reproductive isolation due to differentiated flowering times.

Our study highlights the importance of the IPT-driven evolution of sex allocation to the coexistence of closely related plants. By using a population dynamics model incorporating coevolution of sex allocation towards FSA, our model proposes a new scenario of eco-evolutionary extinction (i.e. “evolution-mediated competitive exclusion”) of competitors. Also, our results suggested two conditions for evolution-mediated competitive exclusion: In the absence of IRC, evolution-mediated competitive exclusion occurs with asymmetry of IPT, whereas in the presence of IRC, a negative correlation between IPT and IRC results in evolution-mediated competitive exclusion. These results indicate that the combination of different types and strength of species interactions is important to determine the outcomes of eco-evolutionary dynamics.

Further empirical and experimental tests could allow for a comparative approach to reveal the importance of the evolution of sex allocation in species coexistence. Incorporating selfing and other mating system characteristics will help us gain a deeper understanding of coexistence and extinction driven by evolution in natural plant communities.

## Acknowledgement

We thank Dr. Hisashi Ohtsuki for his helpful comments. KM was supported by RIKEN Junior Research Associate Program and Grant-in-Aid for JSPS Research Fellow (JSPS KAKENHI Grant Number JP1226645).

## Supplementary figures

**Figure S1:**
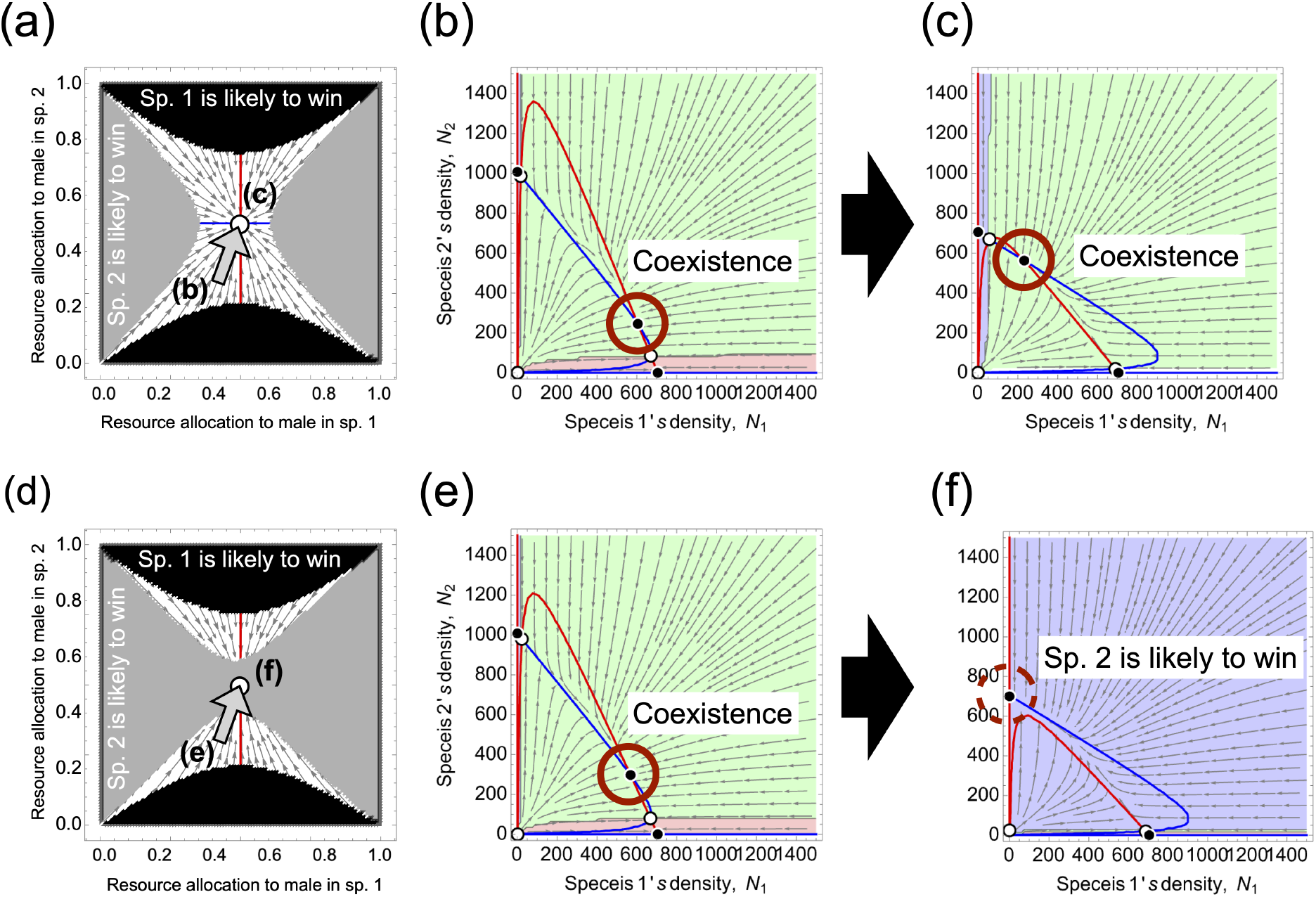
The effects of the difference in initial sex allocation between species 1 and 2 in evolution on coexistence and evolutionary exclusion when IPT is asymmetric and IRC cannot occur. (a, d) Phase diagram of sex allocation in both species. The selection gradient is zero when each species has the FSA (red and blue lines for species 1 and 2, respectively). As the FSA is convergence stable, a species with female-biased sex allocation evolves towards FSA. In contrast, a species already having FSA will not evolve further. Black and gray regions indicate that species 1 and 2 likely win competition, respectively. Gray arrows represent the convergence stability in the evolution of sex allocation. (b, c, d, e) Nullcline of population dynamics of two species. (b, e) When both species have intermediately biased sex allocation, there is a locally stable equilibrium on which coexistence is possible. (c) As the result of evolution towards the FSA in both species, the equilibrium on which coexistence is possible is maintained, but (e) the equilibrium disappears. Green, blue, and red regions indicate the attraction basin of the equilibrium of coexistence and extinction of species 1 and species 2, respectively. Parameters are *α*_21_ = 0.6 (fixed) and (b) *α*_12_ = 0.8, (c) *α*_12_ = 0.9 and the others are in Table 1. Sex allocation is (b, e) 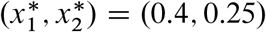, and (c,f) 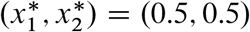.

**Figure S2:**
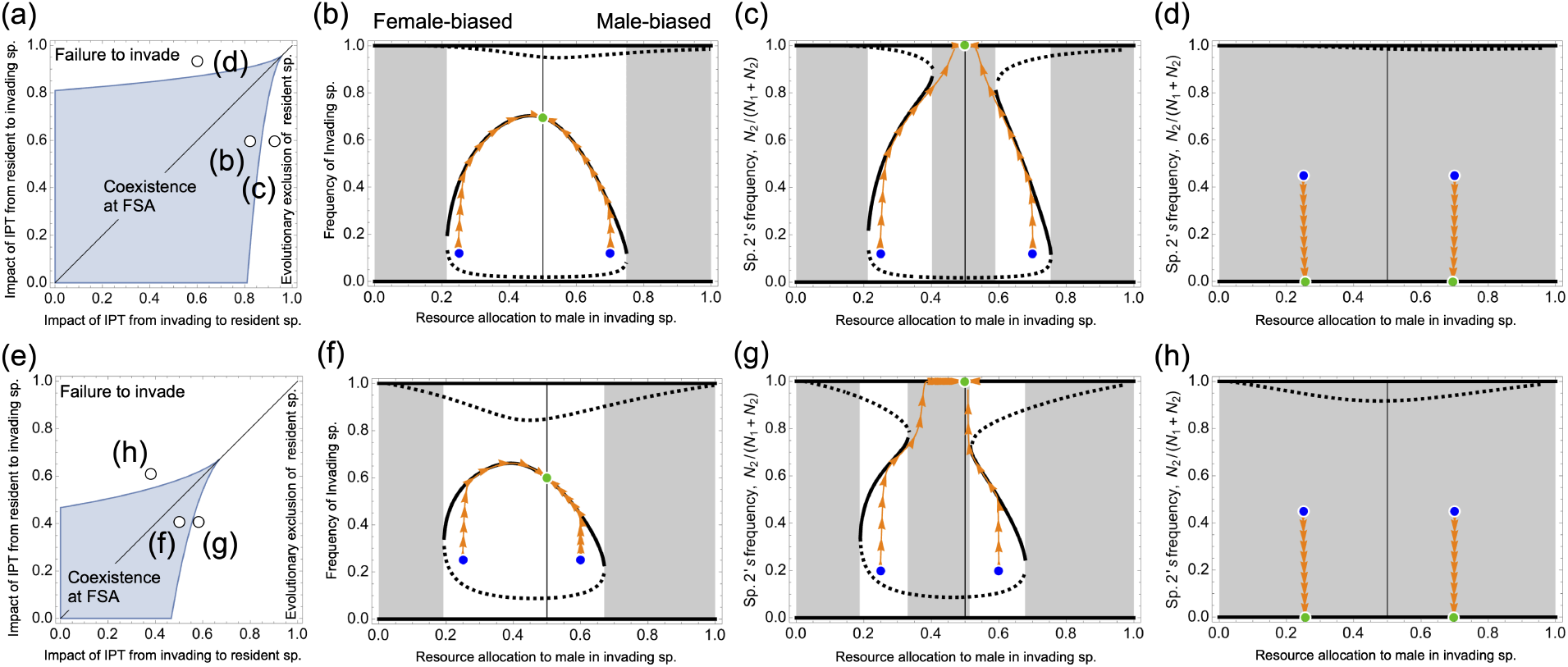
The robustness of evolutionary exclusion in the presence of weak strength of interspecific resource competition between species. (a, d) Parameter dependency of evolutionary exclusion. (b, c, d, f, g, h) Demographic equilibrium along resource allocation to male. Regardless of the strength of resource competition, three outcomes occur. Black dashed and solid lines are unstable and locally stable equilibria. Parameters are (*α*_12_, *α*_21_) = (b, f) (0.8,0.6), (c, g) (0.9,0.6), (d, h) (0.6,0.9) and (a, b, c) *β*= 0 (d, e, f) *β* = 0.1 and the other parameters are in Table 1.

**Figure S3:**
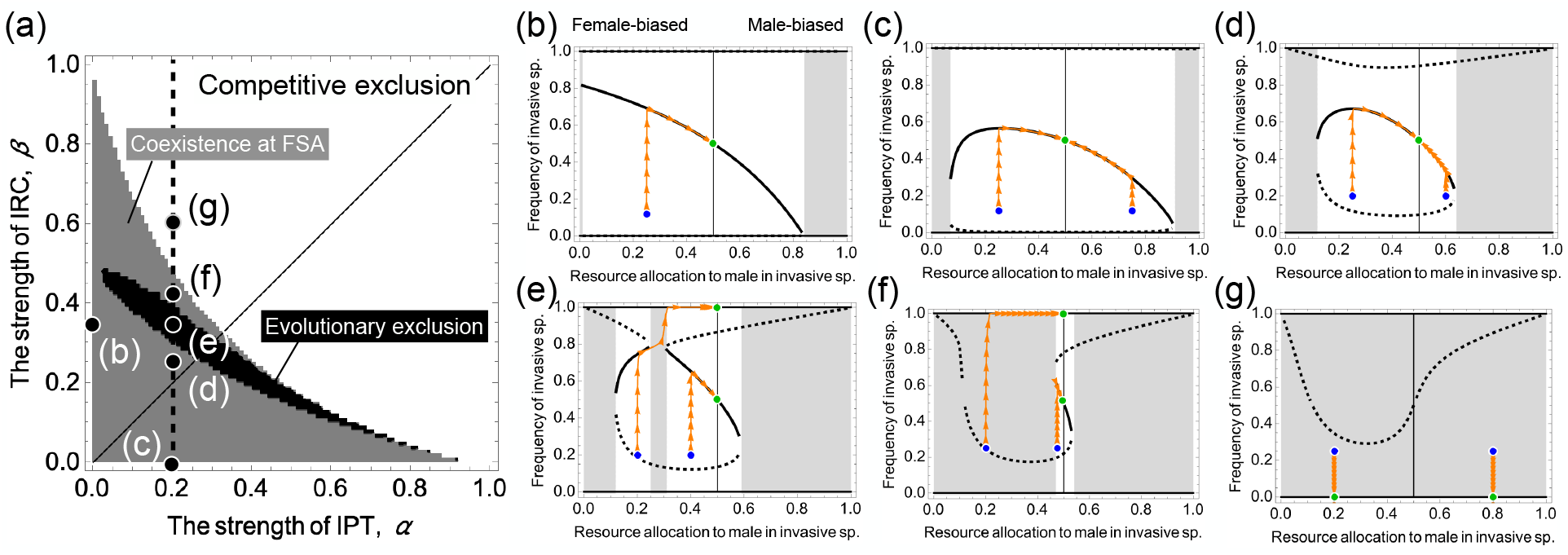
The robustness of evolutionary exclusion in the change in the strength of IPT and IRC. (a) The phase diagram categorizes eco-evolutionary outcomes. (b-g) The relationship between sex allocation and the number and stability of equilibria. Parameters are (*α,β* =(b) (0,0.31), (c) (0.2,0), (b) (0.2,0.25), (c) (0.2,0.31) (d), (0.2,0.4) (e), and (0.2,0.6) and the other parameters are in Table 1.

**Figure S4:**
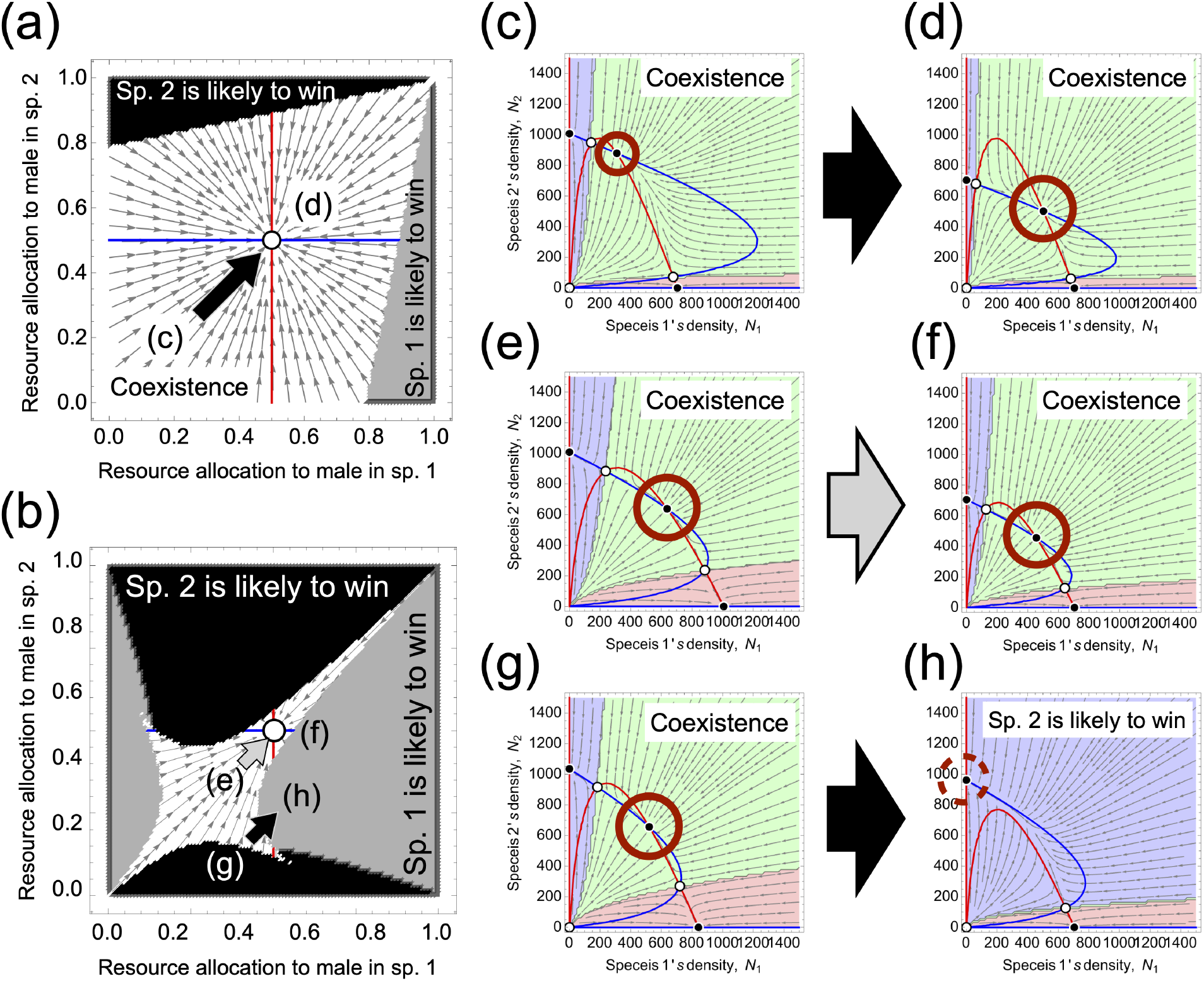
The effects of the difference in initial sex allocation between species 1 and 2 in evolution on coexistence and evolutionary exclusion when IPT is symmetric and IRC is strong. (a, b) Phase diagram of sex allocation in both species. The selection gradient is zero when each species has FSA (red and blue lines). (a) When both species 1 and 2 have female-biased sex allocation, coexistence remains possible. (b) When species 1 has slightly female-biased sex allocation and species 2 has intermediately female-biased sex allocation, the evolution towards the FSA in both species makes species 1 extinct (*i*.*e*., evolutionary exclusion) (black arrow). In contrast, when species 1 and 2 have slightly female-biased sex allocation, such evolution keeps coexistence possible. (c, d, e, f) In the process of evolution towards the FSA in both species, an equilibrium of coexistence is maintained, but (g, h) as the result of evolution towards the FSA in both species, the equilibrium disappears. Parameters are *α* = 0.35, (a, c, d) *α* = *0* and (b, e-h) *α* = 0.2, and the others are described in Table 1. Initial sex allocation is (c, e). 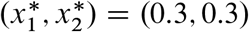, and (g). 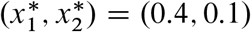.

**Figure S5:**
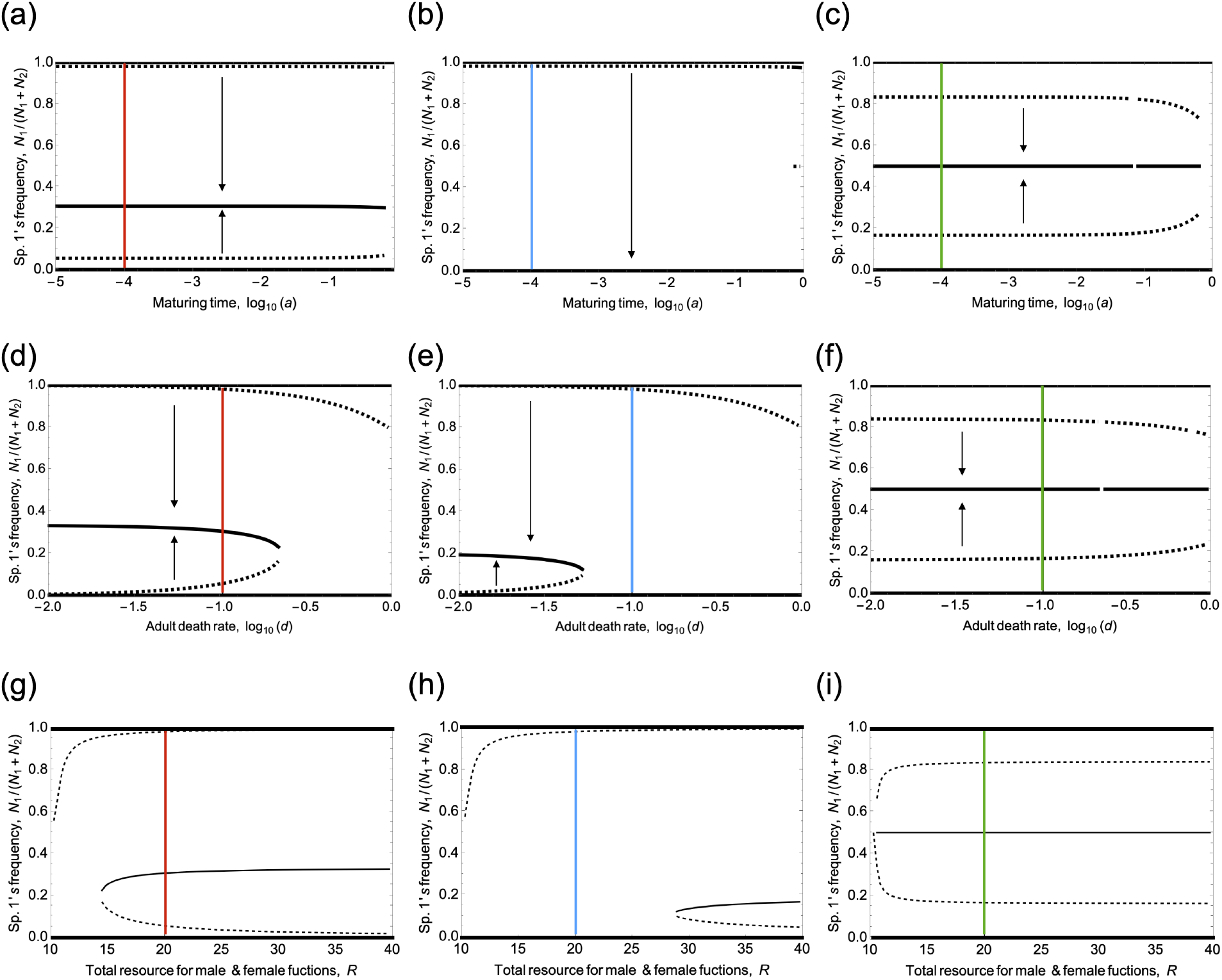
The change in parameters affects the existence of a locally stable equilibrium at the Fisherian sex allocation. Shifts in parameter values cause the change in the number and stability of equilibria if an invading species has the FSA for (a, b, c) the maturing rate of zygotes, *a*_*0*_, for (d, e, f) the adult death rate, *d*, and for (g, h, i) a total resource for the production of pollen grains and ovules, *R*. The left, middle, and right panels show the parameter dependency of results in a case where interspecific pollen transfer is symmetric, but interspecific resource competition cannot occur, in a case where asymmetric interspecific pollen transfer only occurs, and in a case where symmetric interspecific pollen transfer and symmetric interspecific resource competition occurs. Colored lines indicate parameters corresponding to each case. Parameters in each case are described in Table 1.

## Supplementary information

We model the life cycle (birth and reproduction) of a hermaphrodite plant species without selfing. In reproduction, we incorporate pollen transfer within and between species. We show the modeling in Section 1 and derive equations of population dynamics including sex allocation in Section 2 and 3. Furthermore, we derive the invasion fitness from the equations in Section 4. Finally, we derive the condition for which coexistence is possible at the equal sex allocation in both species in Section 5.

### S1. Four processes in a life cycle in plants

The models explicitly incorporate the dynamics of resource competition (Fig. S6a), pollen export and reception within and between species, and reduced mating success due to IPT (Fig. S6b, c). We express male and female costs through the following process: Pollen grains are lost by being exported to heterospecific ovules (Fig. S6b), and ovules are lost by being fertilized by heterospecific pollen grains (Fig. S6c). We consider zygotes to be produced only when a pollen grain successfully mates with a conspecific ovule (Fig. S6d). A model overview is described in Fig. S6.

**Figure S6.**
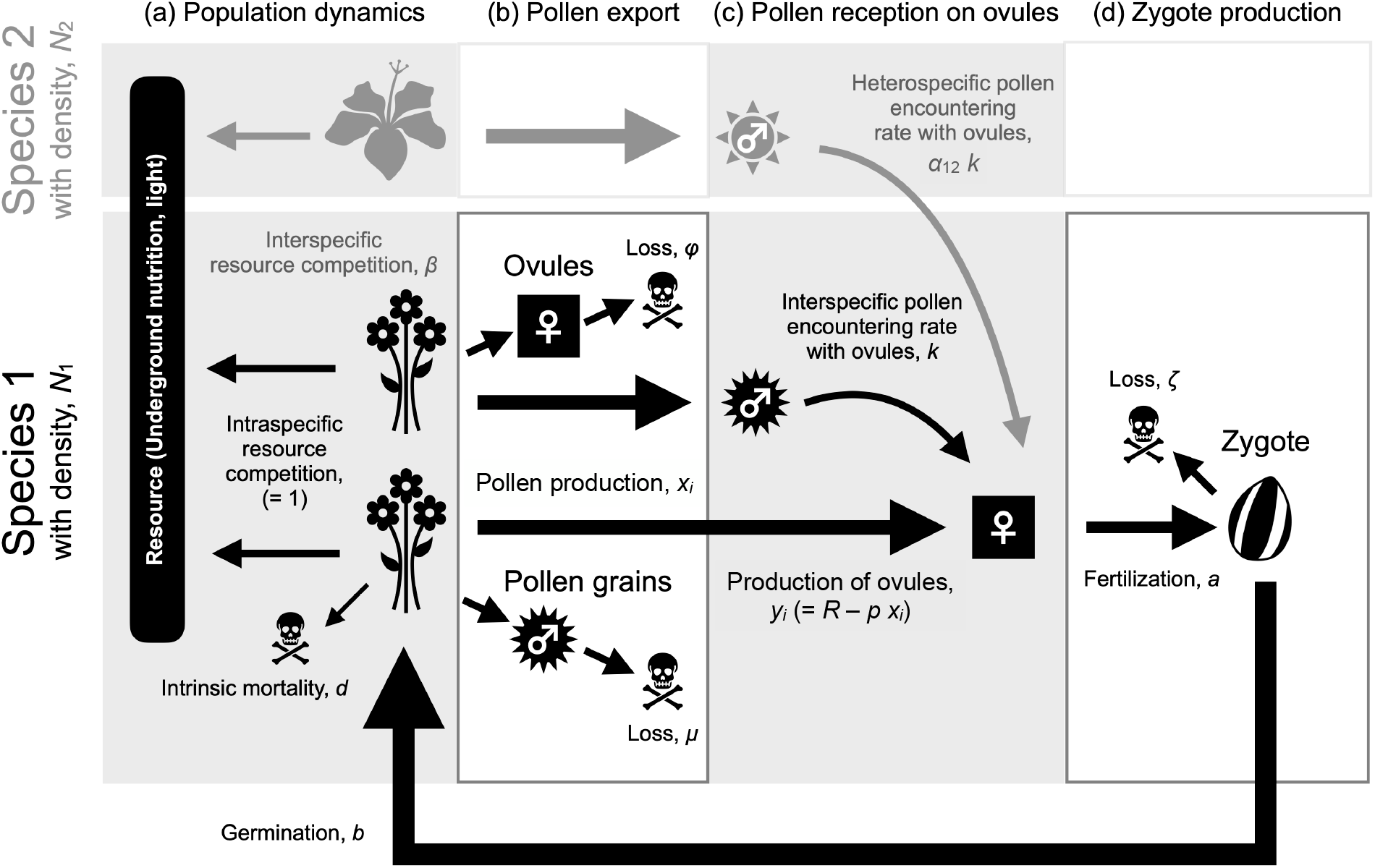
Model overview.

#### a. Population dynamics of adult flowers

We consider two closely related plants that co-flower throughout a year with the density of species *i, N*_*i*_ (*i* = *1; 2*). For simplicity, we consider that one plant individual has one flower. They interact through resource competition and interspecific pollen transfer (IPT). In population dynamics of the flowers with density-dependent mortality rate, *d* (Kishi and Nakazawa 2013; Kuno 1992), two species germinate from seeds with the rate, *b*. Also, flowers may die due to intra- or inter-specific competitions (at a rate, 1 or *β*, respectively). These processes are expressed as:

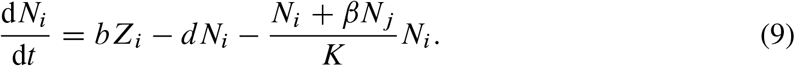

#### b. Production and export of pollen grains

The production rate of pollen grains of species *i* is written as *m*_*i*_. *x*_*i*_ *N*_*i*_ is a total pollen production and *x*_*i*_ is per capita pollen production for species *i*. They are lost at a rate, *µ*, for instance, due to death or transfer to the third plant species. Pollination vectors carry a pollen grain or pollinium to ovules over and over. Pollen grains encounter conspecific and heterospecific ovules depositing *c* conspecific and *h* heterospecific pollen grains with a rate *k*, with the rate in the counterpart species *j* weighed by *α*_*ij*_. These processes are written as:

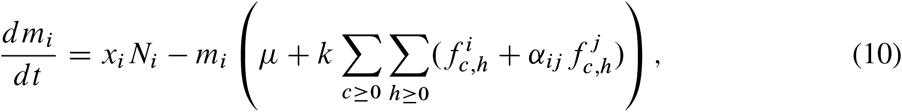

where 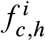 indicates the number of species *i* ‘s ovules depositing *c* conspecific and *h* heterospecific pollen grains. We assume that pollen grains encounter ovules that deposit various numbers of conspecific and heterospecific pollen grains before encountering the pollen grain, including ovules with no pollen grains.

**Figure S7:**
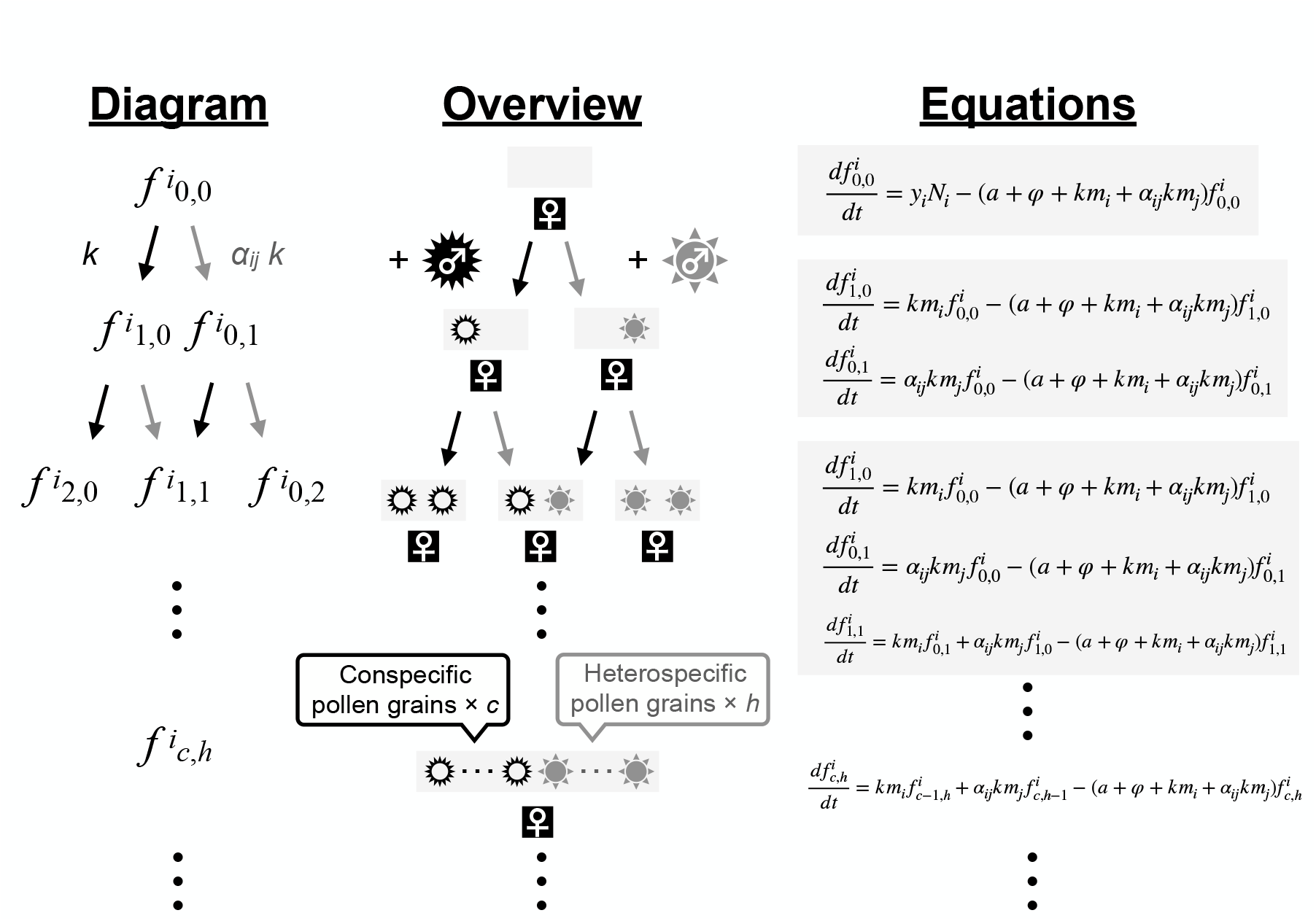
The transition of pollen reception in ovules.

#### c. Production of ovules and pollen reception in ovules

We consider the following transient processes of the number of ovules: Ovules that have received *c* conspecific and *h* heterospecific pollen grains originate either from: (i) ovules with *c* - *1* conspecific and *h* heterospecific pollen grains receive a conspecific pollen grain (which occurs at a rate *km*_*i*_ for species *i*), or (ii) ovules with *c* conspecific and *h* - *1* heterospecific pollen grains receive a heterospecific pollen grain (which occurs at a rate *α*_*ij*_ *km*_*j*_ for species *i*). A flower is closed at a rate *a*, potentially forming a zygote provided that its ovule has received at least one conspecific pollen grain (otherwise, it produces no offspring). The death rate of the ovules (which is common across all types of ovules) is denoted by*φ*. These processes are written as:

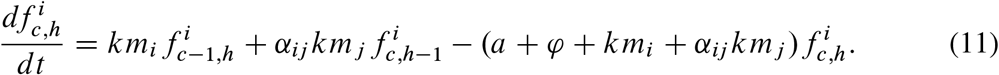

We write the initial state of the process of pollen reception as:

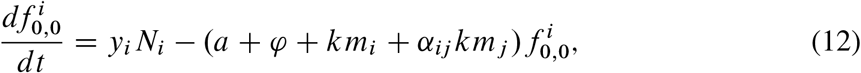

where *y*_*i*_ *N*_*i*_ is a total production of ovules and *y*_*i*_ is per capita production of ovules for species *i*. In Fig. S7, we visualize these processes of pollen reception and we count the number of conspecific and heterospecific pollen grains received by ovules.

#### d. Zygote formation

We write the number of zygotes of species *i* as *Z*_*i*_. We consider that zygotes are produced with the fertilization rate *a*, when an ovule succeeds in mating with one conspecific pollen grain with the probability, *p*_*c,h*_(= *c/*(*c* + *h*)) (i.e. frequency-dependent fertilization). Zygotes become *ζ* mature with a rate *b*, but die at a rate *t*. The dynamics of zygotes is thus given by:

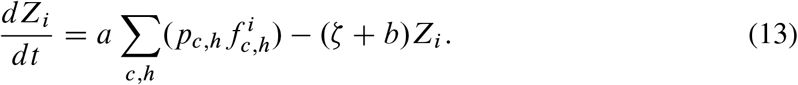

All variables and parameters are defined in Table 2.

### S2. Approximation of time-scale separation in the mating dynamics

Let us assume that the typical timescale of the mating dynamics (i.e. production and export of pollen grains, *dm*_*i*_ */dt*, production of ovules and pollen reception on ovules, 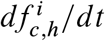, and zygote formation, *dZ*_*i*_ */dt*) is much shorter than the timescale of the population dynamics, *dN*_*i*_ */dt*. That is we assume:

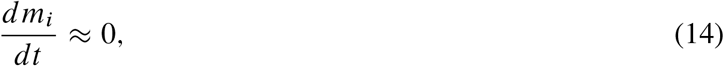

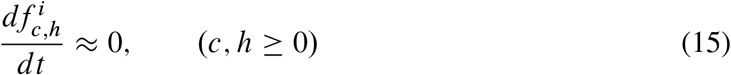

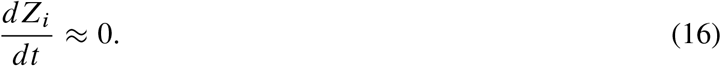

#### a. Pollen dynamics

We assume that the loss rate of pollen grains is much greater than the encountering rate with ovules, indicating no male cost of pollination (i.e. *µ* » *k*). Biologically, we assume that there are many pollen grains. Under the assumption, we solve Eq. (14) as follows:

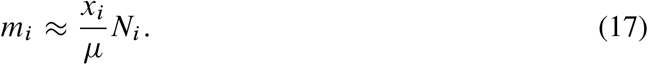

#### b. State transition of what number of pollen grains ovules deposit

Using the time-scale separation of Eq. (15), we get:

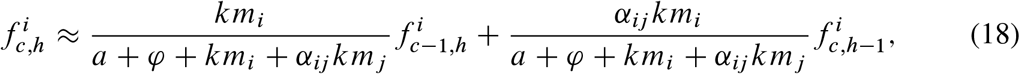

with

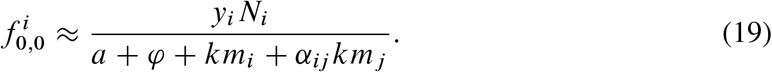

Here, the number of pollen grains on a ovule depends on previous state. By solving Eq. (18, 19) recursively, we get:

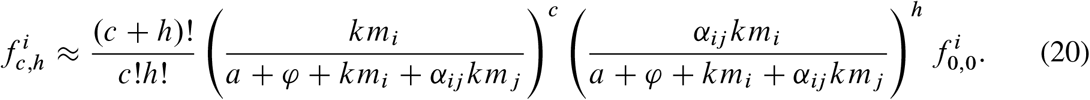

The fraction *km*_*i*_ */*(*a* +*φ*+*km*_*i*_ + *α*_*ij*_ *km*_*j*_) is the probability that an ovule receives a conspecific pollen grain, and the other fraction is the corresponding probability of receiving a heterospecific pollen grain. There are the (*c*+*h*) !/c !*h*! ways of receiving *c* and *h* pollen grains from conspecific and heterospecific flowers (respectively), and thus this formula is interpreted as the binomial process.

#### c. Zygote dynamics

From Eq. (16), we get:

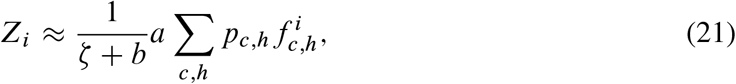

We incorporate *p*_*c,h*_ = *c/*(*c* + *h*) and Eq. (20) into the above equation.

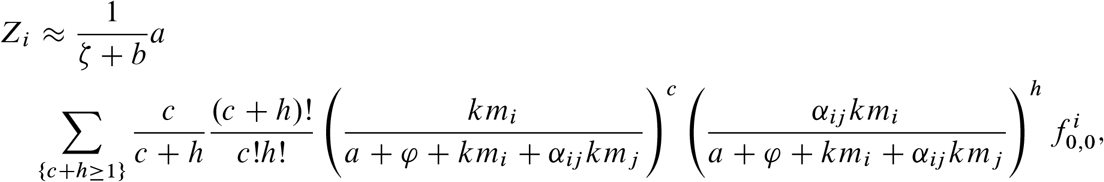

Replacing *c* - *1* = *c ′* and *c* - *1* + *h* = *T* gives:

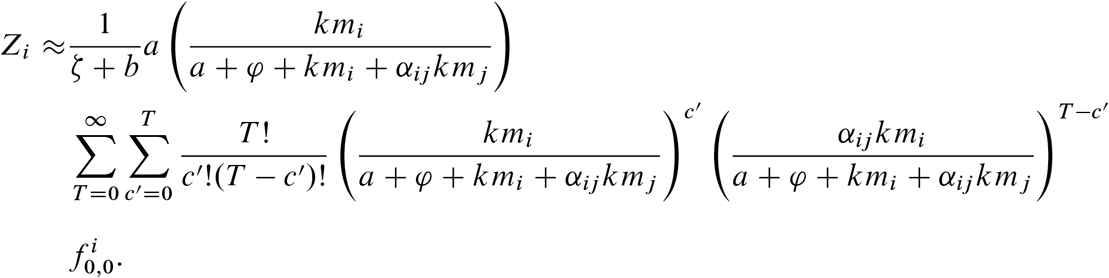

By solving this, we get:

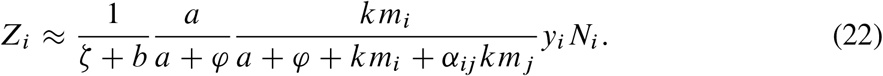

### S3. Derivation of the population dynamics equation including sex allocation

We derive a population dynamics equation including sex allocation by incorporating Eq. (22) of zygote dynamics into the Eq. (9) of population dynamics.

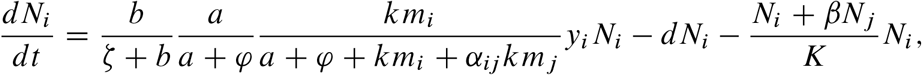

By incorporating Eq. (17) of pollen dynamics into the above equation,

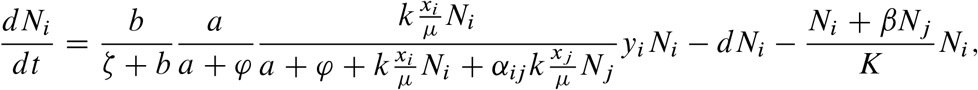

We define the probability of seed germination, *g*_1_ = *b/*(*ζ* + *b*) and fertilization of ovules and conspecific pollen grains, *g*_2_ = *a/*(*a* + *φ*). We write *g* = *g*_1_*g*_2_ for the probability of successful germination. Also, we define probability of ovules’ encountering with conspecific pollen grains as 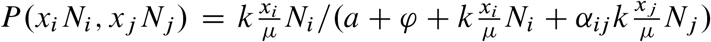. We can therefore rewrite the population dynamics more concisely as:

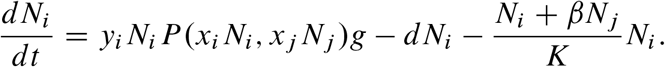

We convert *1=K* to *δ* to get:

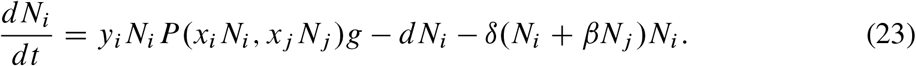

### S4. Adaptive dynamics

#### a. Invasion fitness

To investigate co-evolutionary dynamics of pollen production of two species (*x*_1_, *x*_2_)), we use adaptive dynamics theory (Doebeli 2011; Doebeli and Dieckmann 2000; Hofbauer and Sigmund 1990; Metz, Nisbet, and Geritz 1992). In adaptive dynamics theory, we consider that a monomorphic population of a wild type (Eq. (23)) reaches demographic equilibrium, 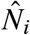 and 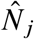 with given phenotypes, *x*_*i*_. Then, evolution occurs when rare mutants with a new phenotype invade the population of a wild type, expressed as the condition where the invasion fitness of mutants with phenotype, 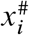 is positive.

The population dynamics of the rare mutants with the density in species *i, n*_*i*_ is written as:

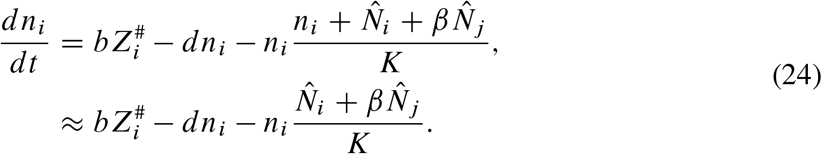

where 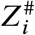 indicates the number of zygotes of mutants, with 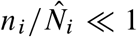 and 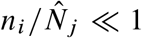.

For the dynamics of male gamete, 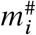 we get:

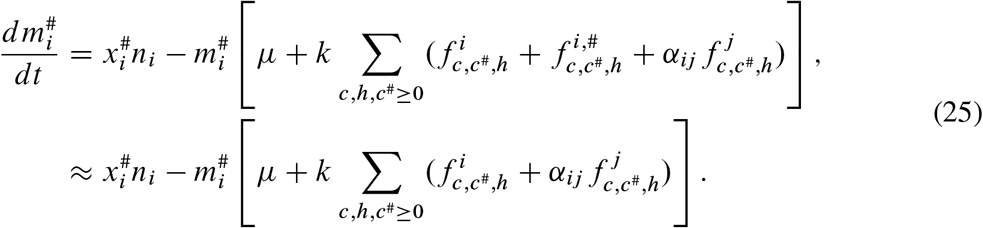

Here, we omit the possibility of male gametes to be received by mutant female gametes (no selfing).

Female compartments are more complicated. We first consider male mating success of the wild type and mutants via encountering with ovules of the wild type. Assuming that (i) mutant pollen does not sire mutant ovules (no selfing), and (ii) a single ovule receives at most single mutant pollen grain (because of the rairity of the mutant), we can write down the ordinary differential equations of the number of ovules sired by a mutant pollen grain as:

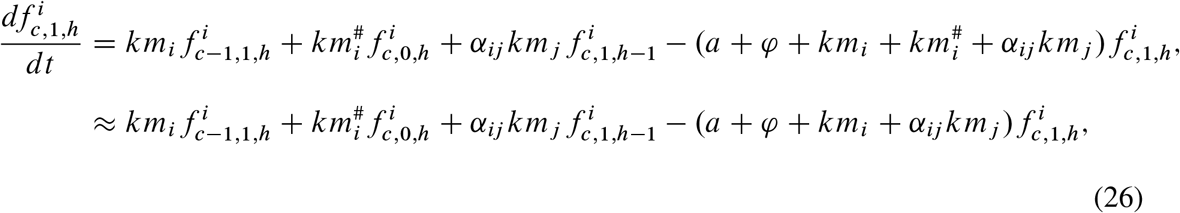

with:

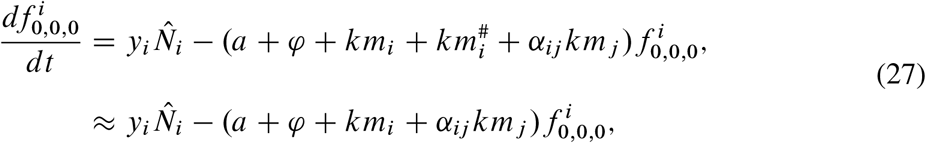

Second, female may receive a mutant pollen grain. The number of fertilized ovules produced by mutants is given by:

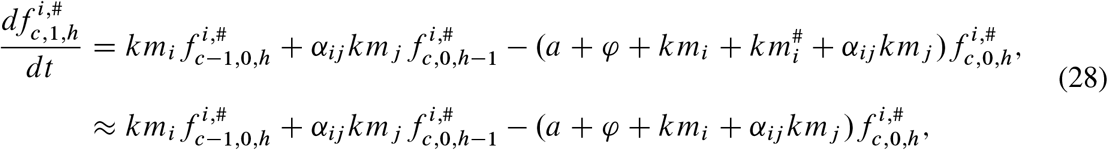

with:

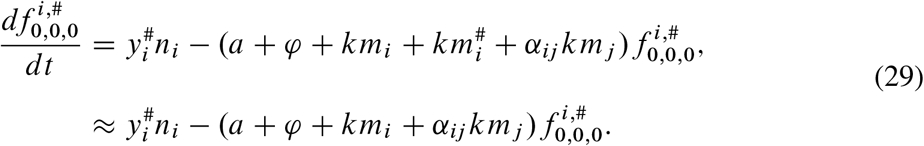

For diploids with no selfing, the number of zygotes that possess the mutant allele is given by:

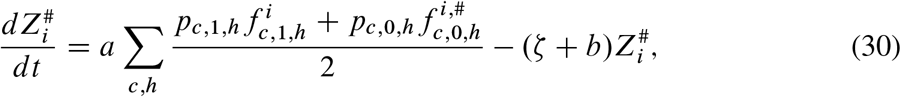

where

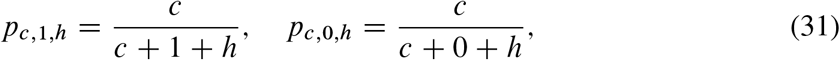

We apply the quasi-equilibrium approximation to the dynamics including mutant phenotype, and solve recursively as in Section 3. The pollen reception, including mutant pollen grains in ovules, is expressed in the following Fig. S8. Colored boxes represent the number of ovules to which no mutant pollen grain is attached.

**Figure S8:**
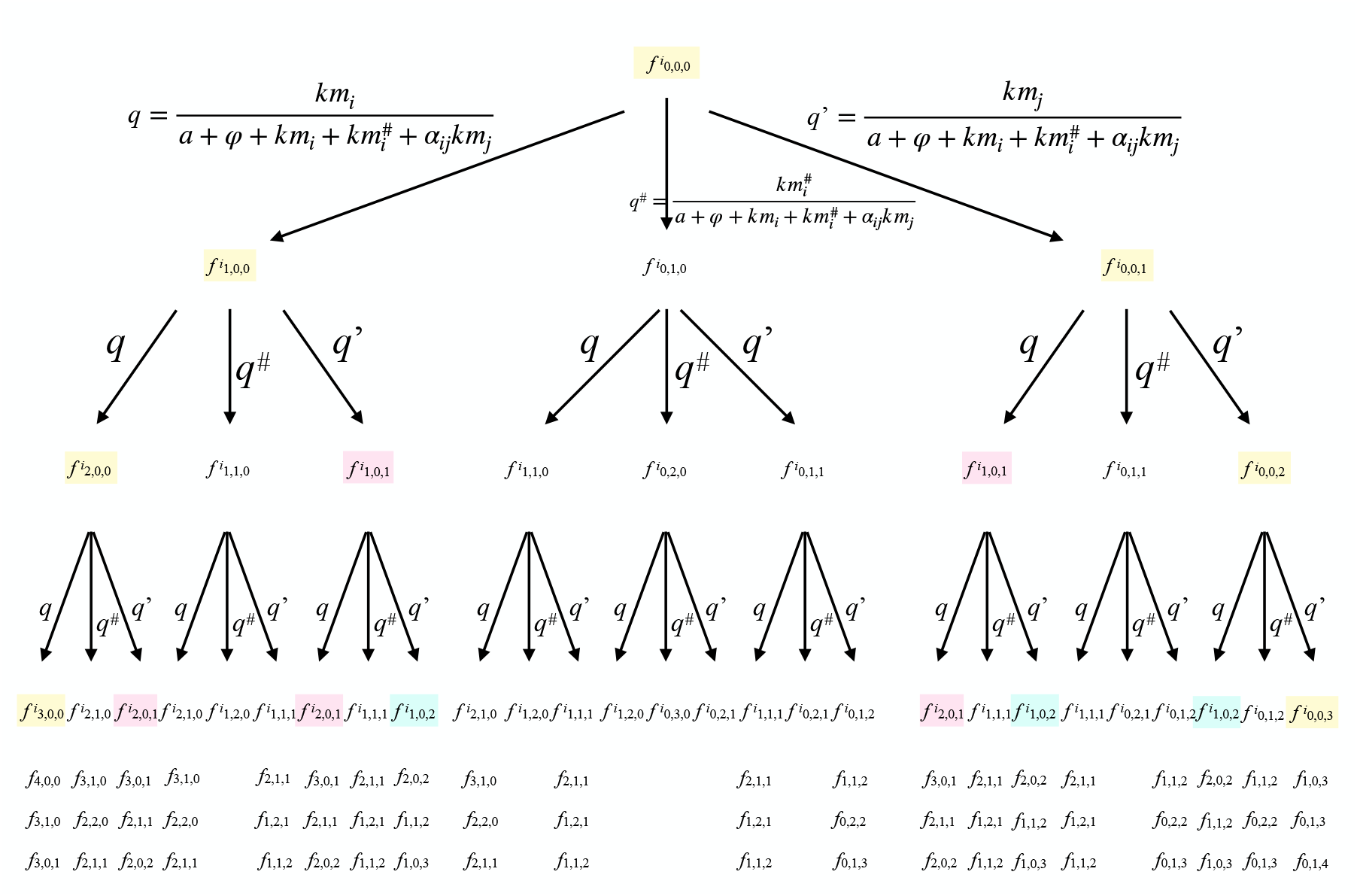
The transition of pollen reception of resident and mutant pollen grains in an ovule.

Thus, we write the number of mutant zygote at equilibrium as:

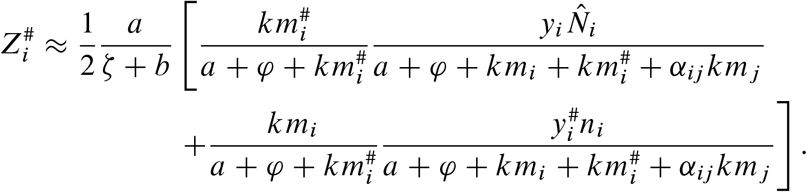

We now use the above number of mutant zygotes to population dynamics of Eq. (24). In addition, considering time-separation approximation in the dynamics of mutant pollen grains in Eq. (25) in a similar way as in Section 2, we get:

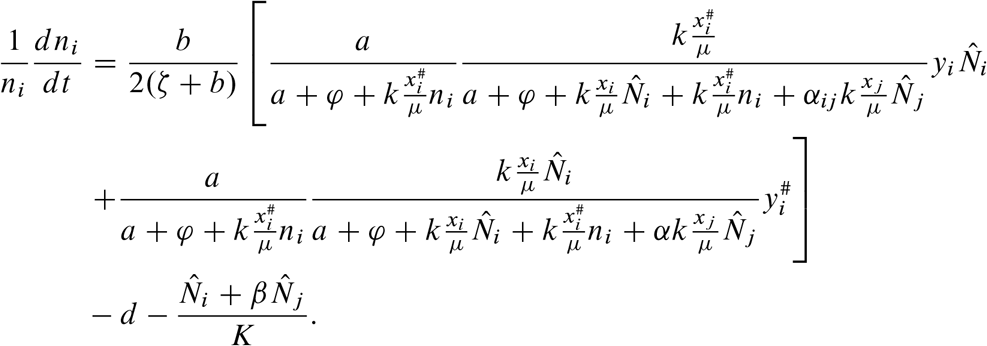

As is similar to Section 3, we convert *1/K* to *δ* to get:

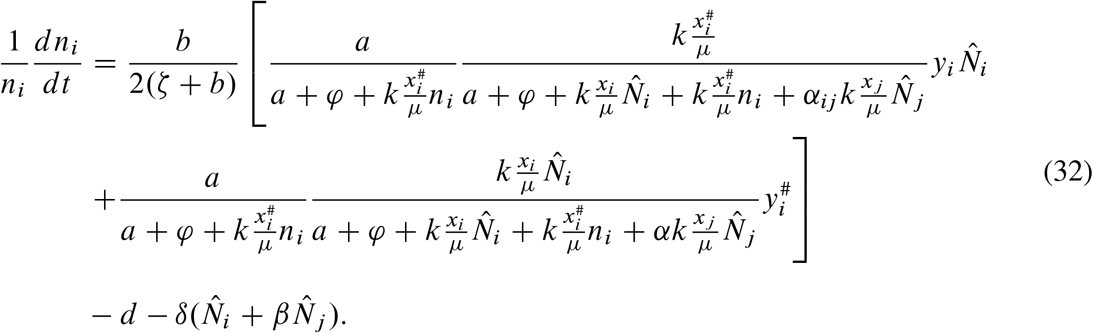

Eventually, assuming that mutants are rare (i.e. *n*_*i*_ ⟶ *0*), the invasion fitness of mutants, 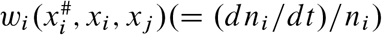 is calculated as:

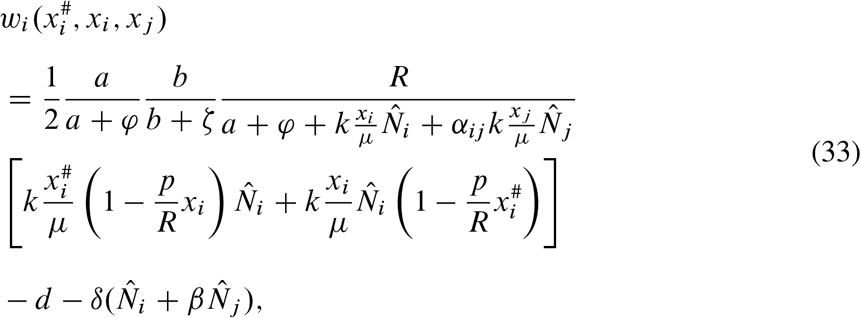

where we assume that a trade-off between resource allocation of male and female functions like *px*_*i*_ + *y*_*i*_ = *R*. For brevity, in the main text, we assume *p* = *1*. The invasion fitness comprises male and female reproductive successes.

Now we write:

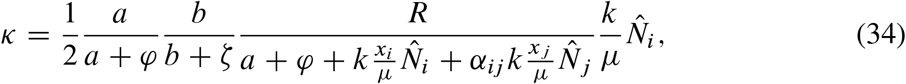

to get:

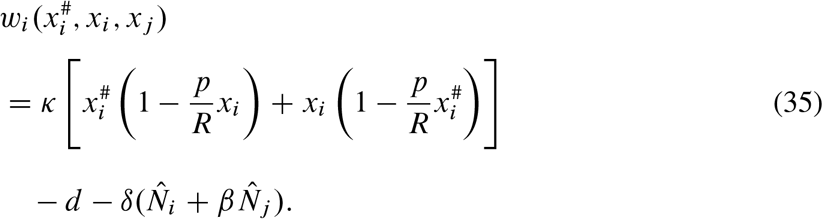

To investigate the direction of evolution, we calculate the selection gradient, *D*_*i*_ (*x*_*i*_, *x*_*j*_) by partially differentiating the invasion fitness of the mutant of species *i*, to get:

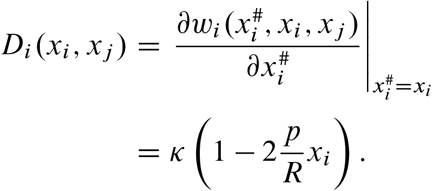

Solving *D*_1_(*x*_1_, *x*_2_) = *D*_2_*(x*_2_; *x*_1_) = 0 for *x*_*1*_ and *x*_*2*_ leads to an equilibrium 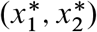 of evolution (i.e. evolutionarily singular point). In the coevolutionary system, we find whether or not evolution converges to the singular point by examining the maximum real parts of the eigenvalues of the following Jaccobean matrix (Dieckmann and Law 1996; Leimar 2009):

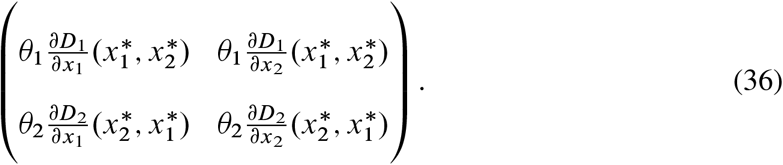

where *θ*_*i*_ tunes the speed of evolution that is determined by mutation rate and population density of each species at equilibrium. When all the real parts are negative, the biological system may reache the singular point (i.e. “convergence stable”). At the singular point 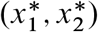, we get the following eigenvalues:

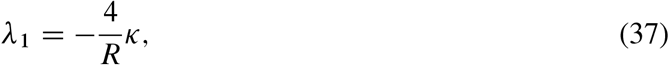

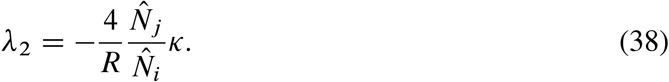

We assume that *K* and *R* are positive constants, and 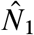 and 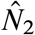 are positive because coevolution occurs when adaptive dynamics theory assumes that two species co-occur. Therefore, eigen-values (37, 38) are always negative and real. Thus, the singular point is always ‘convergence stable.’

We examine whether coexistence is possible in variable sex allocation values of two species, *x*_1_ and *x*_2_ for the cases where IPT occurs in the absence of IRC. In the main text, we have assumed that the resident species has equal sex allocation (FSA) when the invasive species immigrates. Essentially the qualitatively same results follow as we have shown in Fig. 2 even when the initial sex allocation of resident species deviates from FSA (Fig. S1a for case (i) symmetric IPT in which the IPT from species *i* to *j* is equally strong as that from *j* to *i* (*α*_12_ = *α*_21_) and S1d for case (ii) where IPT from an invasive species 2 is much stronger than IPT from a resident species 1 (*α*_12_ » *α*_21_)).

When IPT is symmetric, if an invasive and resident species have intermediately biased sex allocation, coexistence is possible (white region in Fig. S1a). Hence, in the process of evolution towards the FSA in both species, coexistence remains possible (gray arrow in Fig. S1a). A nullcline analysis shows that in this process, a demographic equilibrium of coexistence always exists (Fig. S1b, c). If the invasive species has biased sex allocation and the resident species has extremely female-biased sex allocation, the resident species is driven to extinction (black region in Fig. S1a). In the case where the strength of IPT from the invasive species is much stronger than IPT from the resident species, if the invasive and resident species have intermediately biased sex allocation, coexistence is also possible. Evolution towards the FSA in both species results in evolution-mediated competitive exclusion (gray arrow in Fig. S1d). Again, a nullcline analysis shows that such evolution makes a demographic equilibrium of coexistence disappear (Fig. S1e, f).

In addition, we investigate the robustness of results against changes in the initial values of sex allocation in the presence of IRC. First, we consider a case in the absence of IPT. Our analysis shows that coexistence remains possible in an evolutionary process except for a case where both species initially have extremely male-biased sex allocation before evolution (a white region in Fig. S4a). In this case, nullcline analysis shows that the equilibrium of coexistence is maintained in the process of evolution (a black arrow in Fig. S4a, Fig. S4c, d). In a case where IPT occurs, if both species have slightly female-biased sex allocation, coexistence remains possible in the process of evolution (black arrows in Fig. S4b), whereas if species 2 has extremely female-biased sex allocation and species 1 has slightly female-biased sex allocation, evolution-mediated competitive exclusion occurs (a gray arrow in Fig. S4b). A nullcline analysis shows that a locally stable equilibrium of coexistence always emerges in the process of evolution (Fig. S4e, f), while the equilibrium disappears as a result of evolution-mediated competitive exclusion (Fig. S4g, h). Sex allocation in both invasive and resident species evolves to FSA independently with each other and regardless of ecological conditions including IPT. Conversely, the ecological dynamics of these species depend on both species’ sex allocation. As a result, the eco-evolutionary outcome is determined by how the evolutionary trajectory moves through the regions of qualitatively different ecological consequences (the white, gray, and black regions in Fig. S1a and S1d).

The robustness arises because the regions and sequence of evolutionary trajectories remain unchanged, whether or not the resident species’ sex allocation is restricted to FSA.

### S5. Coexistence at equal sex allocation and extinct boundary in a phase diagram of co-evolutionary dynamics

In the absence of IRC, when coexistence is possible at intermediately biased sex allocation but either species goes extinct at the Fisherian sex allocation (FSA), evolution-mediated competitive exclusion occurs. We examine the condition for which either species goes extinct at the FSA. Also, in the presence of IRC, when coexistence is possible at extremely biased sex allocation but either species goes extinct at intermediately biased sex allocation, evolution-mediated competitive exclusion may occur, even if coexistence is possible at the FSA. We examine whether the equilibrium of coexistence exists at sex allocation which ranges from *0* to the FSA. For brevity, we carry out a conventional variable-transform so that otherwise two-species dynamics can be converted to the dynamics of a ratio of densities of two species: *ρ* = *N*_*1*_*/N*_*2*_ (Otto and Day 2007).

First we derive the population dynamics including the allocation to male and female functions, as follows:

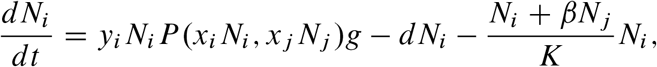

where 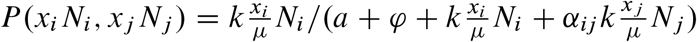

Now, we define *a′* = *a* + *φ* and *B* = *k/µ*, to get:

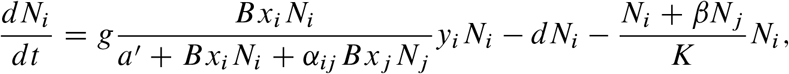

Assuming *a′* « 1 and *φ < a*, we get:

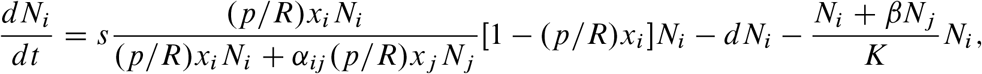

where *s* = *gR*.

Because *p=R* is constant, we rewrite *X*_*i*_ = (*p/R*)*x*_*i*_, to get:

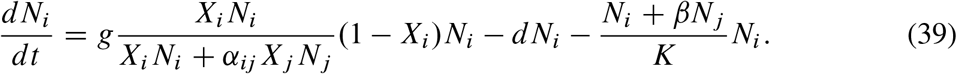

At demographic equilibrium (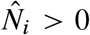 (for *i =* 1,2)), population growth rate is zero, so we get:

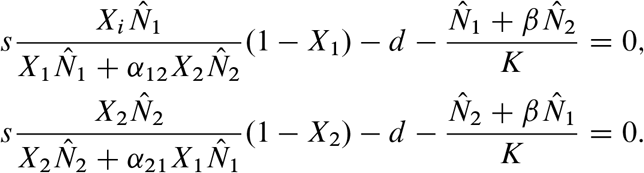

Expressing 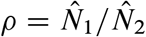, we get:

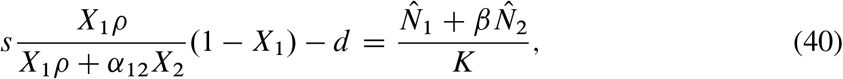

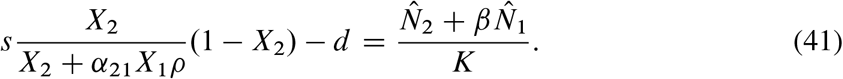

By dividing equation (40) by equation (41) given that *X*_2_ ≠1, we get:

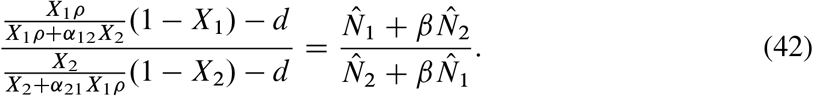

#### a. Asymmetric strength of IPT in the absence of IRC

We assume no interspecific resource competition (IRC, i.e. *β* = 0), in which case we get:

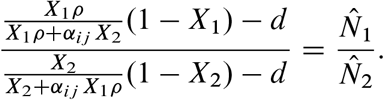

That is,

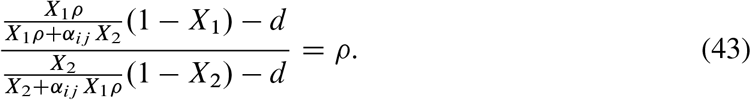

We investigate the condition for which evolution-mediated competitive exclusion occurs. Evolution-mediated competitive exclusion occurs in the process of evolution from biased sex allocation at which coexistence is possible towards equal allocation at which either species are extinct. That is, we examine the condition for which coexistence is possible at FSA (*X*_1_ = *X*_2_ = 0.5). Inserting *X*_1_ = *X*_2_ = 0.5, we can obtain the condition as:

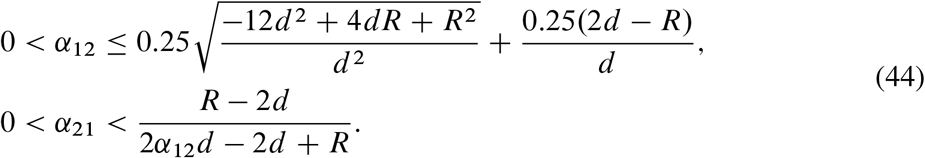

or

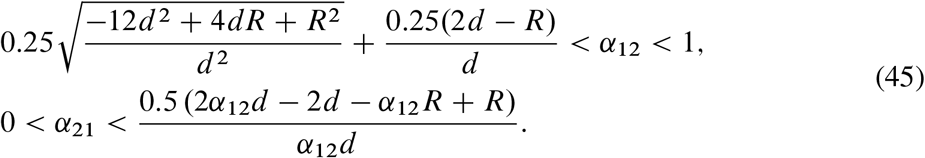

For example, at *R* = 20, *d* = 0.1, we get supplementary Fig. S2a.

#### b. Symmetric strength of IPT in the presence of IRC

For simplicity, we assume symmetry of IPT (i.e. *α*_12_ = *α*_21_ = *α*) in the presence of symmetric IRC (i.e. *β >* 0). In this condition, we replace 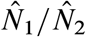 with *ρ*, to get:

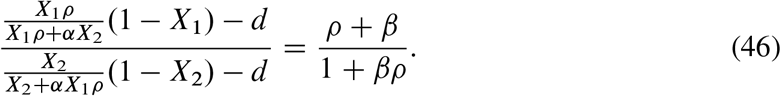

We rewrite the equation above to the following cubic polynomial:

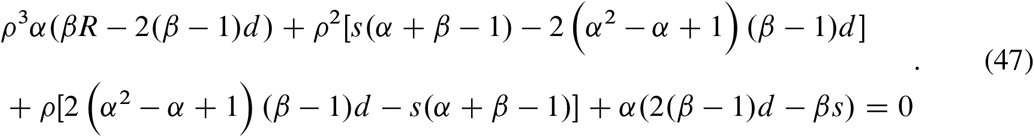

First, we investigate the condition for which coexistence is possible at FSA, as is similar to the above section (a). We consider a relationship among the three real roots (Ψ_1_, Ψ_2_, and Ψ_3_) and coefficients in the cubic polynomials:

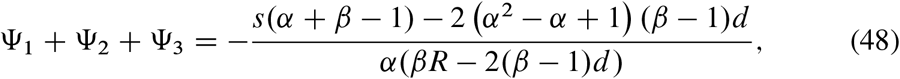

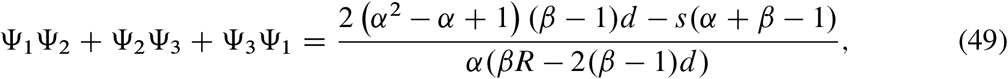

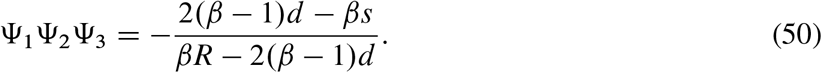

Because equation (50) is always positive, we examine the signs of equations (48) and (49). For example, at *R* = *20, d* = *0:1*, we obtain the condition for which coexistence is possible at FSA (the gray area in the supplementary Fig. S3a).

Next, we investigate the condition for evolution-mediated competitive exclusion, given that one species has equal sex allocation. We focus on the transitions between a case where one species is driven to extinction and a case where coexistence is possible. That is, we derive the condition for which equation (47) has two real roots on *X*_*i*_ when *X*_*i*_ ranges from *0* to *0:5* provided that *X*_*j*_ is 0.5 (for *i, j* = 1, 2). For example, at *R* = 20, *d* = 0.1, we obtain the condition for which there are two regions where coexistence is impossible (the gray area in the supplementary Fig. S3a) when *X*_2_ ranges from 0 to 0.5 (the black area in the supplementary Fig. S3a).

We examine that ecological outcomes are affected by the parameters other than the strengths of IPT and IRC. We first relax the assumption under which the half saturation pollen density in the absence of competing species is negligible (*a*_0_ = 0), by considering *a*_0_ *>* 0. Also, Eq. (45, 46, 49-51) show that the coexistence condition depends on a total resource to produce pollen grains and ovules, *R*, and natural mortality of adults, *d*. Changing the values of the three parameters alters the occurrence of a locally stable equilibrium where coexistence is possible at the FSA. For *a*_*0*_, regardless of the strengths of IPT and IRC, a wide range of the parameter values produces the same number of the locally stable equilibrium (Fig. S5a, b, c). For *d* and *R*, when IPT is symmetric in the absence of IRC, the number of the locally stable equilibrium is the same in a small value of *d* (Fig. S5d) and in a large value of *R* (Fig. S5g), while the number is the same when IPT from the invasive species is strong in the absence of IRC in a large value of *d* (Fig. S5e) and in a small value of *R* (Fig. S5h). In a case where IPT is symmetric and IRC is strong, a wide range of the parameter values produces the same number of the equilibrium (Fig. S5f, i).

## Supplementary tables

**Table 2:** Symbols and Their Definitions on the Supplementary information.

| Symbols | Definition | Default values (if any) |
| --- | --- | --- |
| $i$ or $j$ | Species label | - |
| $t$ | Time | $0 \leq t$ |
| $N_i$ | Density of species $i$ | Dynamic variables |
| $R$ | The total resource a plant invests in the production of pollen grains and ovules | $0 \leq R$ |
| $p$ | The resource allocation necessary to produce a pollen grain relative to that of an ovule | 1 |
| $x_i$ | Pollen production of a plant | Evolutionary trait |
| $y_i$ | Ovule production of a plant | $R - px_i$ |
| $a$ | Fertilization rate of ovules | 0.001 |
| $\varphi$ | Loss of ovules | $a/10$ |
| $a_0$ | Half saturation pollen density in the absence of competing species | $a + \varphi$ |
| $b$ | Birth rate of adult flowers | 1 |
| $\zeta$ | Loss of zygotes | $b/10$ |
| $g$ | Germination fraction | $(a/(a + \varphi))(b/(b + \zeta))$ |
| $d$ | Natural mortality rate | 0.1 |
| $\mu$ | Loss of pollen grains | 0.1 |
| $k$ | Encountering probability of ovules with heterospecific pollen grains | 0.001 |
| $\alpha_{ij}$ | The rate at which a heterospecific pollen grain is transferred to an ovule (relative to that of a conspecific pollen grain) | $0 \leq \alpha_{ij} \leq 1$ |
| $K$ | Carrying capacity of adult flowers | 100 |
| $\delta$ | Coefficient of density-dependent mortality | $1/K$ |
| $\beta$ | The strength of interspecific resource competition relative to that of intraspecific competition | $0 \leq \beta \leq 1$ |

